# Fatty acid metabolite promotes lymphatic identity in stem cell-derived endothelial cells

**DOI:** 10.1101/2025.11.03.686405

**Authors:** Donghyun Paul Jeong, Sanjoy Saha, Daniel Montes-Pinzon, Angela Taglione, N. Keilany Lightsey, Rananjaya Subash Gamage, Brendan Stein, James Brandon Dixon, Donny Hanjaya-Putra

**Author notes:** Address correspondence to: Donny Hanjaya-Putra, 2010G McCourtney Hall East University of Notre Dame Notre Dame, IN 46556.

## Abstract

The lymphatic system plays various crucial but underappreciated roles in fluid transport and immune response in numerous organs and tissue types. Consequently, generation of lymphatic vessels has been postulated as an innovative therapeutic strategy for various diseases. However, there is a lack of efficient and reliable method to differentiate human pluripotent stem cells into lymphatic endothelial cells (LEC) for lymphatic regeneration. Current differentiation methods suffer from poor yield and low lymphatic marker expression, while also having limited clinical applicability due to its reliance on either the embryoid body intermediates or xenogenic supporting cells. Given that LECs exclusively rely on anaerobic and fatty acid metabolism due to the hypoxic environment of the lymph, here we report that the unique lymphatic-specific metabolic pathways can be exploited to promote lymphatic identity in differentiated LECs (dLECs). We show that dLECs express elevated levels of lymphatic markers compared to native endothelial cells, which is up to 15 times higher than the current leading standard of dLECs. Moreover, dLECs can form lymphatic vascular networks in both 2D and 3D, as well as secrete important lymphangiocrine for organ maturation. Upon implantation into double-ligation tail lymphedema and mammary fat pad models, dLECs were able to integrate with the host lymphatic vessels, restore fluid flow, and reduce swelling. Collectively, we show that metabolite supplementation can drive stem cell differentiation into dLECs, which can be incorporated into new alternative methods for personalized therapies and disease modeling, as well as provide a direct therapeutic option for lymphedema and lymphatic disorders.

**Graphical Abstract:** 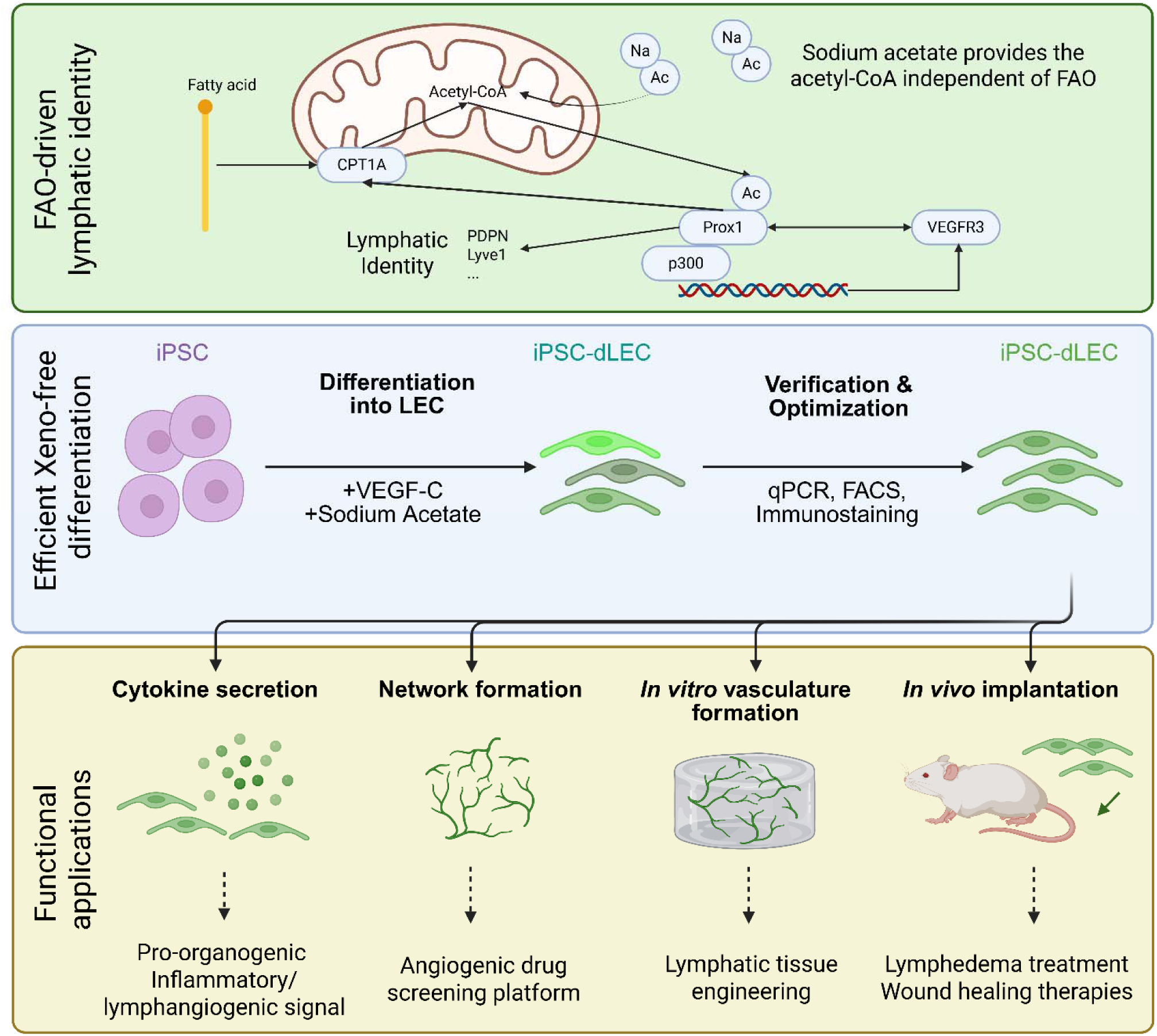

A xeno-free and stepwise differentiation protocol with VEGF-C and sodium acetate induces lymphatic identity in iPSC-derived endothelial cells. Differentiated LECs (dLECs) express key lymphatic markers as verified using bulk RNA-sequencing, qPCR, FACS, and immunostaining. These dLECs secrete important cytokines for lympangiocrine signaling, able to form robust 2D and 3D lymphatic networks, as well as demonstrate *in vivo* functionality and host integration in murine models.

## Introduction

The lymphatic system is a secondary circulatory system that works in conjunction with the blood vessels to maintain fluid homeostasis, lipid transport, and immune trafficking.^1–3^ The lymphatic system consists of lymph nodes, lymphoid organs, and lymphatic capillary vessels and ducts, which collect the fluid coming from capillary filtration from the blood vessels and return them back into circulation.^4,5^ Despite their importance, the lymphatic system is not as well-understood compared to its blood counterpart, in part due to a relatively small list of lymphatic markers that distinguish it from the blood circulatory system.^6^ In a circulatory system including the lymphatics, endothelial cells (ECs) constitute the inner lining of vessels and therefore serve as the direct barrier between the fluid, either blood or lymph, and the tissue.^7^ Therefore, ECs play a key role in regulating vessel permeability, constriction, and flow rate, and at times even plays an endocrinal role by producing cytokines that are important for organ maintenance and maturation.^8–11^ Likewise, lymphatic endothelial cells (LECs) play a similarly important role in regulating lymphatic performance. Proper network formation of LECs is important for preventing swelling that is commonly associated with lymphatic diseases such as lymphedema.^12,13^ LECs also secrete lymphangiocrines such as reelin which play a role in cardiac tissue maturation and other organ development.^14,15^

Because of their importance, LECs derived from human induced pluripotent stem cells (hiPSCs) have been a focus of research in the field of lymphatics. These human differentiated (dLECs) have therapeutic potential as they can promote lymphangiogenesis for wound healing and lymphatic remodeling for lymphedema patients.^16,17^ They are also crucial for developing hiPSCs-derived engineered tissues that are physiologically relevant as part of the new alternative methods (NAMs) for drug screening and disease modeling^18–20^ However, existing studies on hiPSCs-derived endothelial cells have mostly focused on blood vasculature system, mainly due to the difficulty of specifically targeting LECs which are similar to ECs in biomolecular and physiological characteristics. Existing differentiation protocols mostly rely on the formation of embryoid body intermediates or xenogenic supporting cells (i.e., mouse stromal cells, OP9), which suffer from low differentiation efficiency, reduced viability, and low clinical applicability due to xenogenic cells.^21–23^

Further complicating the efforts to develop hiPSCs derived dLECs is the lack of definitive theory on the origin of LECs during embryonic development. The centrifugal theory posits that the lymphatic vessels develop by sprouting from venous endothelium, whereas the centripetal theory suggests that the lymphatic vessels are formed from the mesenchymal cells undergoing *in situ* differentiation.^24^ Human iPSCs, given their analogous nature to embryonic cells, can be differentiated to target mature cells by mimicking the embryogenic environments associated with the target cell line.^25–27^ Therefore, it is intuitive that differentiation of hiPSCs to LECs should also follow the embryogenic origins of LECs. There have been some evidence that transdifferentiation from blood endothelial cells (BECs) to LECs is possible, and other papers have identified pathways that regulate the LEC-BEC separation.^28,29^ However, a recent study into EC specification during mouse embryogenesis discovered that Prox1 and LYVE-1 positive LECs are derived from angioblasts rather than venous ECs, suggesting that a viable pathway of LECs differentiation directly from hiPSCs through angioblast stage is possible.^30^

In this study, we focus on unique metabolic characteristics of LECs to further drive the differentiating ECs to lymphatic phenotype. Previous approaches have relied mainly on LEC-specific growth factors such as vascular endothelial growth factor-C (VEGF-C), which is a ligand to LEC-specific vascular endothelial growth factor receptor-3 (VEGFR3).^21,31,32^ In addition to growth factors, we focus on the LEC’s unique dependence on fatty acid metabolism for its energy needs and gene expression. Native LECs are exposed to highly hypoxic environment of the lymph, resulting in the overreliance of LEC on anaerobic metabolism compared to its blood counterparts.^33^ In addition, fatty acid oxidation (FAO) produces acetyl-CoA, which is used for histone acetylation of VEGFR3, a lymphatic marker that has a positive feedback loop with Prox1, the master lymphatic regulator gene.^34,35^ Therefore, we explored the use of FAO-related metabolites in addition to growth factors to promote lymphatic identity expression in hiPSC-derived endothelial cells.

## Results

### Sodium acetate treatment results in expression of key lymphatic markers

To verify our hypothesis that unique lymphatic metabolic pathways can be exploited to promote lymphatic identity, we tested sodium acetate and VEGF-C supplementation in stepwise differentiation protocols and compared them to the co-culture differentiation protocol as a control **(Figure 1)**. Most existing growth factor-dependent differentiation protocols rely on the two-or three-stage differentiation process, where the hiPSC are first differentiated into mesodermal stage and then further differentiated into endothelial lineage.^36–39^ We have tested our lymphatic maturation process to both the three-stage **(Figure 1A)** and the two-stage (**Figure 1B**) protocols. In the three-step protocol the hiPSCs are differentiated to (i) mesodermal cells then (ii) endothelial progenitor-like cells, and finally to (iii) lymphatic endothelial cells, whereas in the two-stage protocol the hiPSCs are first driven to (i) mesodermal cells then directly differentiated to (ii) lymphatic endothelial cells. We have found that the three-stage differentiation process yields up to 60 times greater cell population expressing lymphatic identity when supplemented with sodium acetate and VEGF-C (**Supplementary Figure 1**).

**Figure 1.**
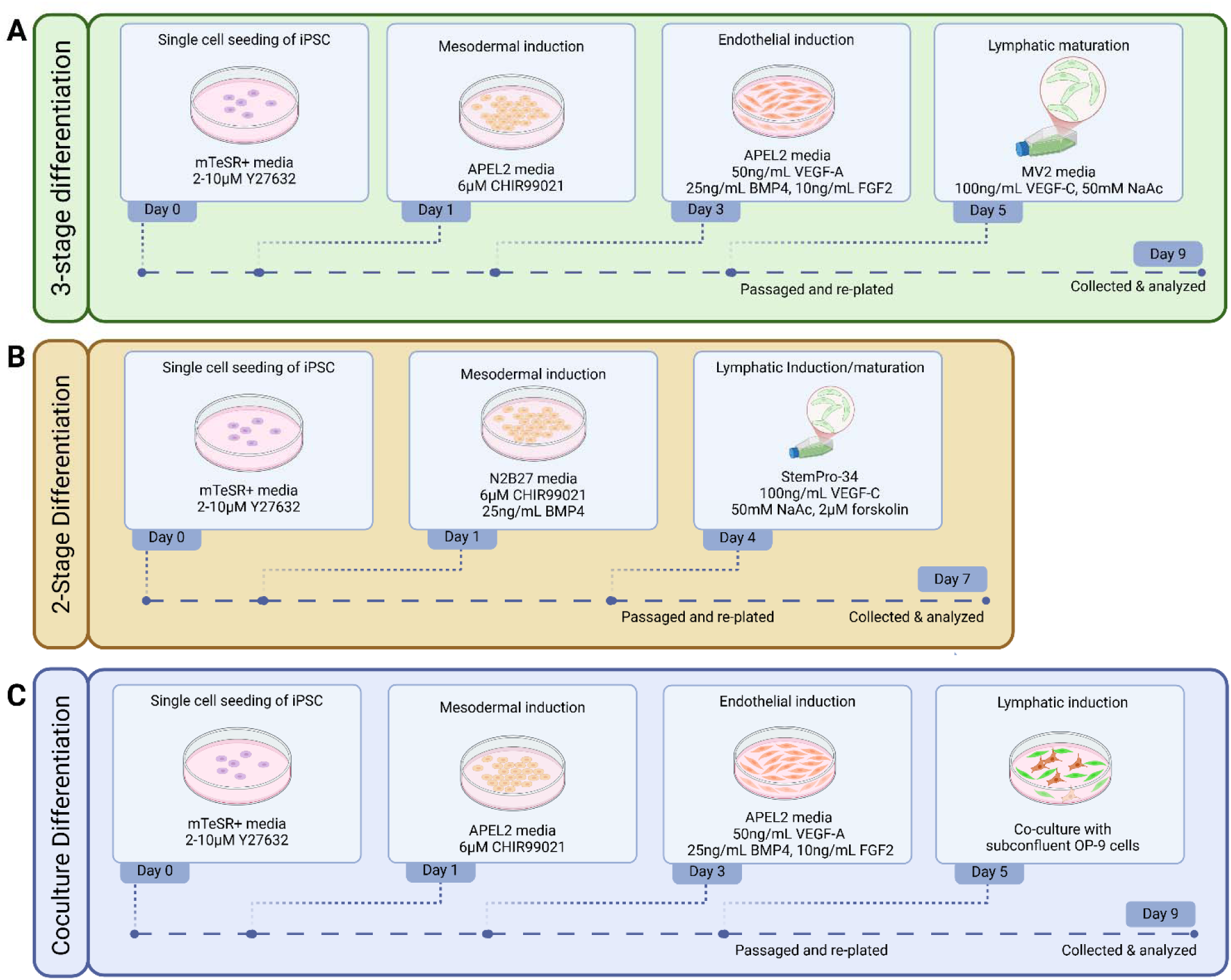
Differentiation Protocols. Summary of the differentiation protocol tested in his study. (**A**) The three-stage differentiation protocol is inspired by the BEC differentiation protocol published by Harding and colleagues, which we have determined to be best suited for inducing lymphatic identity.^36^ The lymphatic maturation stage consists of VEGF-C and sodium acetate treatment for 4 days. (**B**) The two-stage differentiation protocol is adapted from a BEC differentiation protocol published by Patsch and colleagues with a similar lymphatic maturation process.^37^ **(C)** The coculture method is modified from a protocol published by Kono and colleagues.^22^ The three-stage and two-stage protocols were modified to include VEGF-C and sodium acetate during lymphatic maturation phases, and coculture protocol was modified to be compatible with hiPSC differentiation rather than the human embryonic stem cells the original publication used.

In the three-stage differentiation process, to drive hiPSCs into their mesodermal lineage, we first culture hiPSCs as single cells suspension with 10 μM Y27632 in mTeSR+ media for 1 day, followed by 2 day treatment with APEL2 media and 6 μM CHIR99021 GSK inhibitor. This was followed by treatment with 50 ng/mL of VEGF-A, 25 ng/mL of BMP4, and 10 ng/mL of FGF2 in APEL2 media for 2 days **(Figure 1A)**. After that, the differentiated cells were replated at 50,000-80,000 cells/cm^2^ density, which should be optimized for each cell type (**Supplementary Figure 2**). Then the cells are treated with 50 ng/mL of VEGF-C and sodium acetate, a precursor molecule to acetyl-CoA, in microvascular endothelial growth medium (EGM-MV2) media for 4 days (**Figure 2A**).

**Figure 2.**
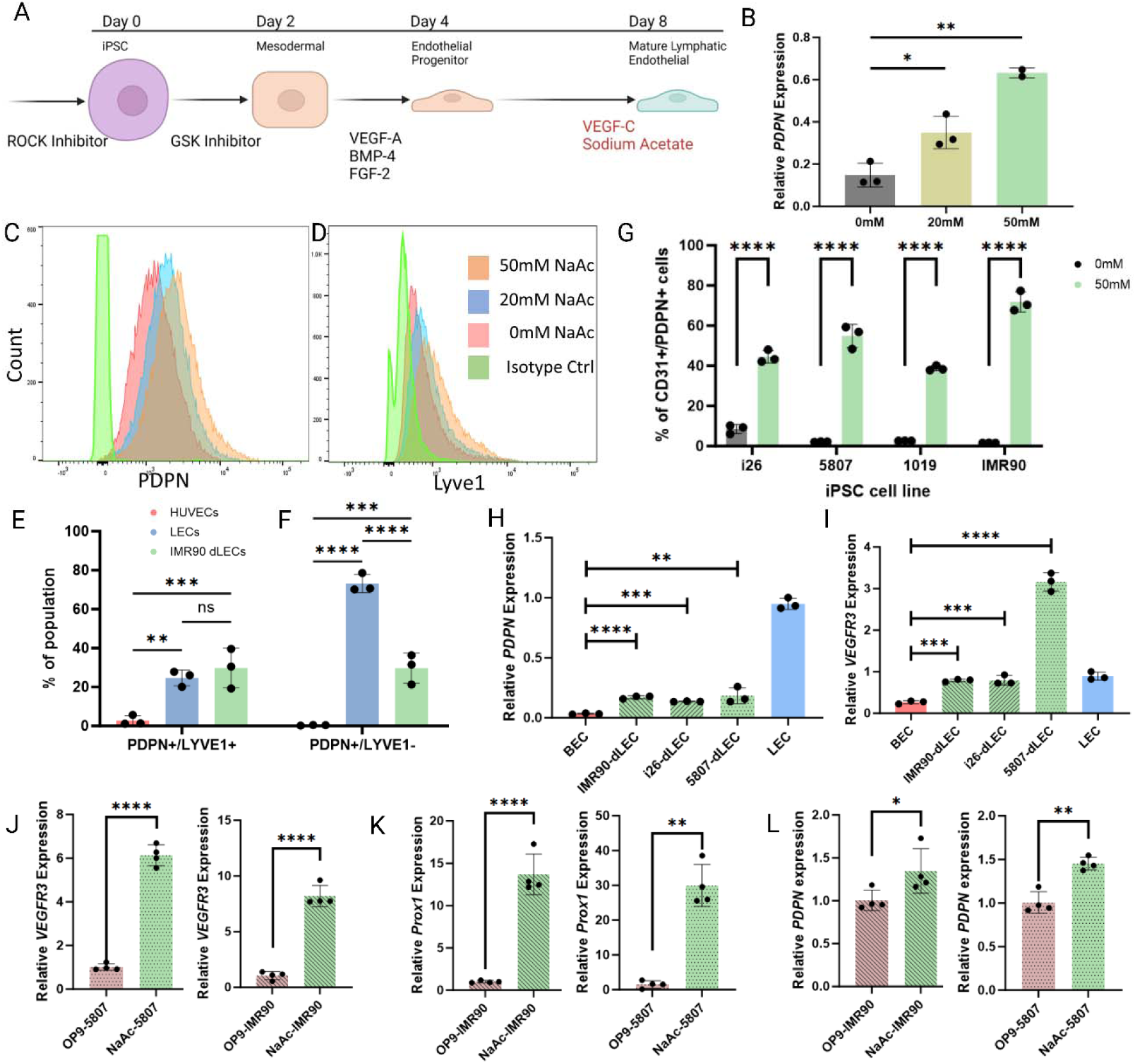
Expression of lymphatic markers with sodium acetate and VEGF-C treatment. **(A**) The differentiation schematic outlining our modifications to existing iPSC-EC differentiation protocol. (**B**) Relative podoplanin-mRNA expression for 0mM, 20mM, and 50mM NaAc treatment groups. The expression is shown as relative to GAPDH expression for each group. (**C-D**) Representative histogram showing shifts in PDPN and LYVE-1 expression as NaAc concentration increases. Isotype control is shown in light green and serves as a basis for threshold for marker-expressing cells (**Supplementary Figure 2**). (**E-F**) Percentage of PDPN+/Lyve1+ double positive population and PDPN+ single positive population for dLECs compared with LECs. (**G**) Percentage of PDPN^+^/CD31^+^ population resulting from our protocol with and without NaAc treatment. For each iPSC cell line, we ran the differentiation protocol in triplicates. (**H**) VEGFR3 expression showing the increase in our differentiated cells compared to native BEC and LEC. (**I**) qPCR results showing an increase in PDPN expression in all three cell lines tested compared to BEC. (**J-L**) qPCR results showing a higher expression of three lymphatic markers in dLECs generated from our metabolite-based method when compared to the standard OP9-based method. *p<0.05, **p<0.01, ***p<0.001, ****p<0.0001. All statistical tests were performed with at least three replicate datapoints using Student’s *t*-test.

We optimized the concentration of sodium acetate added at the last stage and found that 50 mM concentration of sodium acetate (NaAc) in the final differentiation media results in optimal increase in lymphatic markers. We observed a over 2-fold increase in podoplanin (*PDPN*) mRNA expression in cells treated with 20mM NaAc compared to 0 mM treatment, and we observed a over 4-fold increase with 50 mM NaAc treatment (**Figure 2B**). We also observed a similar trend with flow cytometry data analyzing protein level expression of PDPN and LYVE-1, another key lymphatic identity marker (**Figure 2C-D**). We did not use concentrations higher than 50 mM as it resulted in high level of cell mortality, most likely due to the pH shift For all subsequent experiments, we subject the cells to 50 mM NaAc treatment along with 50 ng/mL VEGF-C for the final 4 days.

We observed that our dLECs yield approximately 40% of PDPN+ cells overall, with approximately 20% expressing both Lyve1 and PDPN. (**Figure 2E-F, Supplementary Figure 7**). This closely matches native LECs which also exhibit around 20% of Lyve1/PDPN double positive population. Our protocol resulted in between 80-90% of CD31^+^ cells **(Supplementary Figures 3-6)**. When cells were treated with only VEGF-C and no sodium acetate, the protocol yielded less than 3% of CD31^+^/PDPN^+^ cells. However, with the addition of sodium acetate, the yield of the double-positive cells rose to approximately 40%. To ensure reproducibility of our protocol, we repeated this method with four different hiPSC cell lines derived from various cell types, as well as from male and female donors (**Supplementary Table 1**). We found similar increases in CD31^+^/PDPN^+^ cell yield with sodium acetate treatment (**Figure 2G, Supplementary Figure 1, Supplementary Figure 3-6**).

We also tested for mRNA-level expression changes in *PDPN* and *VEGFR3* and found elevated levels of both LEC markers compared to native BECs, indicating sodium acetate treatment combined with VEGF-C results in expression of key lymphatic markers at the transcriptomic level. We observe a comparable level of *VEGFR3* expression compared to native LECs (**Figure 2H**). Particularly, 5807 hiPSC line demonstrated over 3-fold increase in expression of *VEGFR3* over native LECs. *PDPN*, which is a downstream marker from the VEGFR3-Prox1 feedback loop, also experienced over 5-fold increase in *PDPN* expression over BECs, at approximately 25% expression level of native LECs (**Figure 2I**). Taken together, these results indicate that sodium acetate treatment, when combined with VEGF-C treatment, can directly increase *VEGFR3* transcription which results in subsequent increases in the expression of key lymphatic markers.

### Xeno-free differentiation yields improvement over co-culture method

Currently, the established method for differentiating stem cells into LECs relies on co-culture with a mouse stromal OP9 cell line. Kono and colleagues published a method to differentiate embryonic stem cells into LECs by co-culturing the VEGFR2^+^ cells in a sub-confluent culture of OP9 cells through VEGF-C-dependent pathway.^22^ Similarly, others found that co-culturing hiPSC-derived early endothelial cells with OP9 cell line led to efficient differentiation into LECs.^21,40^ As shown earlier, VEGF-C alone is not sufficient to induce LEC phenotypes in early endothelial cell line (**Figure 2G**), suggesting that OP9 plays a crucial and indispensable role in LEC differentiation. However, the reliance on a mouse cell line introduces potential complications in reproducibility and clinical relevance.

Therefore, we compared our differentiation protocol, which eliminates the need for a mouse stromal OP9 cell line with the additional metabolite treatment along with VEGF-C, to the existing OP9 co-culture standards. We followed the protocol described Kono et al. for two hiPSC cell lines and compared the lymphatic marker expression with that of our dLECs (**Figure 1C**). Our protocol yields over 10-fold increase in *VEGFR3* and *Prox1* expression for both hiPSC lines compared to the OP9 method, indicating that our proposed pathway of inducing the VEGFR3-Prox1 positive feedback loop through sodium acetate treatment is more effective than the current OP9 method (**Figure 2J-K**). In addition, our method also results in over 30% improvement of *PDPN* expression, which is downstream of our proposed feedback loop, indicating that our method not only activates the *Prox1* master regulator gene more effectively, but also results in a higher downstream lymphatic marker expression (**Figure 2L**). Taken together, these results indicate a significant improvement over the current standard of OP9 method, yielding higher lymphatic marker expression while also maintaining the benefits of a xeno-free differentiation method.

### Differentiated LECs express Prox1, the master regulator of lymphatic identity

We hypothesized that the increase in VEGFR3 expression caused by the sodium acetate and VEGF-C treatment would subsequently affect *Prox1* expression, a master regulator of lymphatic identity. We performed our differentiation protocol with the removal of CD144^-^ cells on day 4. Subsequently, we performed immunostaining on native LECs, native BECs, and our dLECs derived from three different iPSC cell lines (**Figure 3A**). We quantified the immunostaining images by selecting region of interest (ROI) based on DAPI stains through Cellpose, then calculating the average fluorescent intensity within each ROI. We observe comparable levels of ERG, a generic endothelial marker expressed in both BEC and LEC (**Figure 3B**). In comparison, we saw a comparable level of Prox1 expression in dLEC compared to LEC, while BEC saw negligible levels of Prox1 (**Figure 3C**).

**Figure 3.**
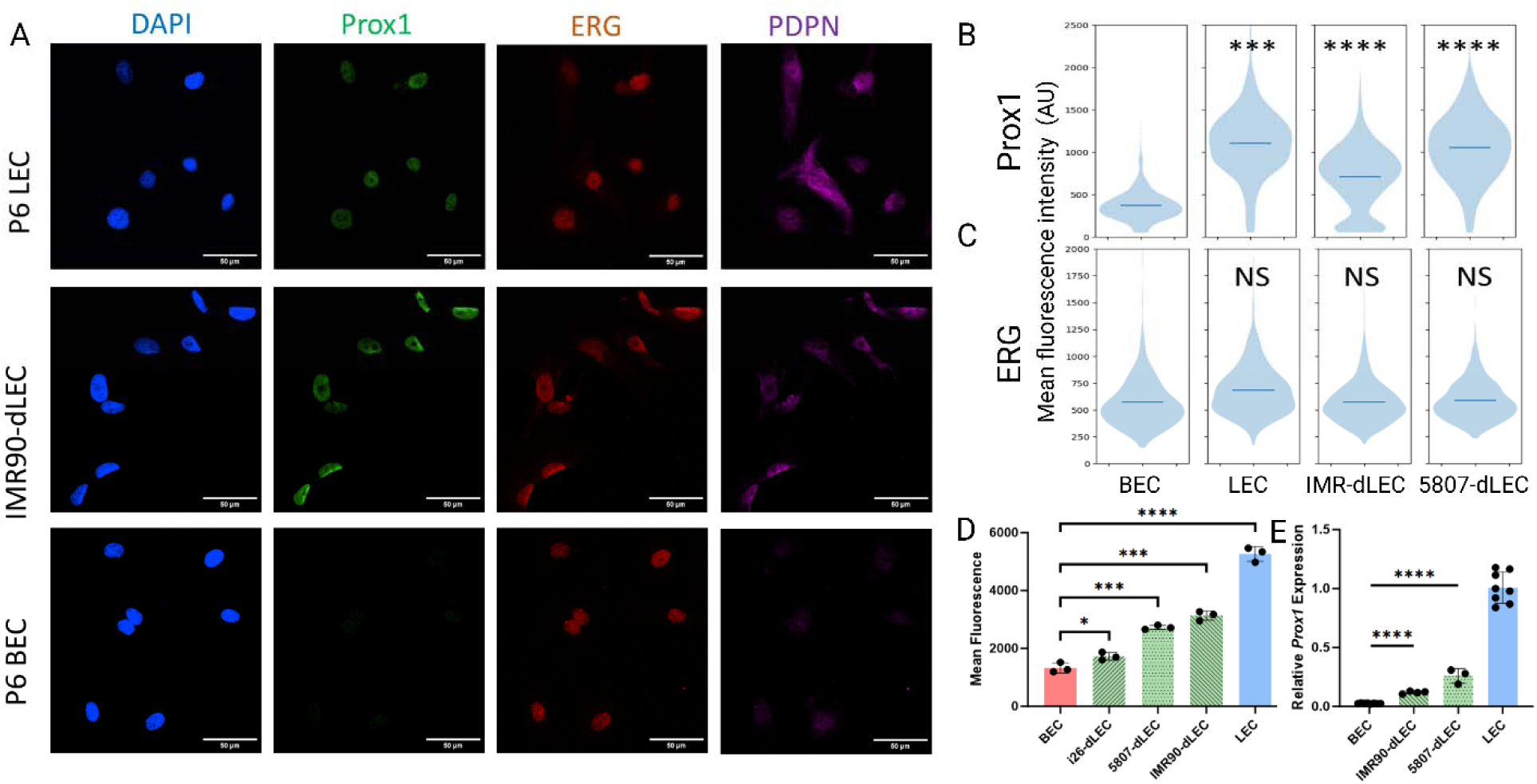
Prox1 expression in the differentiated LECs. (**A**) Immunostaining results comparing the expression of Prox1 and ERG between native LEC, BEC, and dLEC. The cells were fixed, permeabilized, then stained with DAPI, Prox1, ERG, and PDPN. Scale bar represents 50 μm. Representative images are shown. (**B-C**) Quantification of the immunostaining results. The nuclei of each cell were determined based on DAPI staining through Cellpose algorithm, then the average intensity for ERG and Prox1 channels were quantified for each cell. The overall population across all replicates are shown in the violin plot. All statistical significances are relative to the BEC condition. (**D**) Quantification for the FACS results based on the mean fluorescence intensity. Samples were double stained ith ERG and Prox1, and ERG+ cells were selected and quantified for mean Prox1 signal. Each sample was run with 3 separately cultured and stained populations. (**E**) Real-time qPCR results showing an increase in Prox1 expression in all three cell lines tested compared to BEC. *p<0.05, **p<0.01, ***p<0.001, ****p<0.0001. All statistical tests were performed wit at least three replicate datapoints using Student’s t-test.

We further show the expression of Prox1 using flow cytometry data. We observe that our dLECs express comparable levels of Prox1 compared to LEC, and significantly higher than BEC (**Figure 3D**). This Prox1 expression is maintained at a similar level at least up to third passaging as measured by immunostaining data, allowing us to use the dLECs at least up to P3 for our subsequent studies (**Supplementary Figure 8**). Interestingly, the i26 cell line seemed to have a statistically significant but significantly lower Prox1 expression compared to other dLECs and LEC, and more comparable to BECs. The remaining two cell lines that we tested demonstrated Prox1 levels of approximately 50% of that of LECs. The i26 is the only cell line we tested that was PBMC-derived, indicating that fibroblast-derived hiPSC may be better suited for LEC differentiation. We further verified that the increase in Prox1 level is shown in RNA expression as well, although interestingly, the difference between dLEC and LEC in Prox1 RNA expression was more pronounced than that of protein level expression, indicating that there may be subtle differences in Prox1 transcription and translation in our dLECs (**Figure 3E**).

### Differentiated LECs have similar transcriptomic profile to native LECs

In order to ascertain that our dLECs express LEC genes in addition to the identified marker genes, we performed an overall transcriptomic analysis on our cells compared to the native counterparts. We collected the RNA samples from the differentiated cells along with native LEC, BEC, and undifferentiated hiPSC and performed bulk RNA sequencing. We performed alignment analysis using Spliced Transcripts Alignment to a Reference (STAR) RNA-seq aligner and performed dimensionality reduction using principal component analysis on the resulting gene count matrix (**Supplementary Figure 9**). The resulting 2D plot shows our dLECs are distinct from hiPSC and BEC and similar to LECs, as shown by the relative distances between clusters (**Figure 4A**). We selected top 1% of genes with highest variability in expression level between LEC and hiPSC and plotted the Z-score for each sample type for each gene, and found that our dLECs are distinct from hiPSC and similar to LECs, while BECs have a distinct differential expression for several genes including (**Figure 4B**).

**Figure 4.**
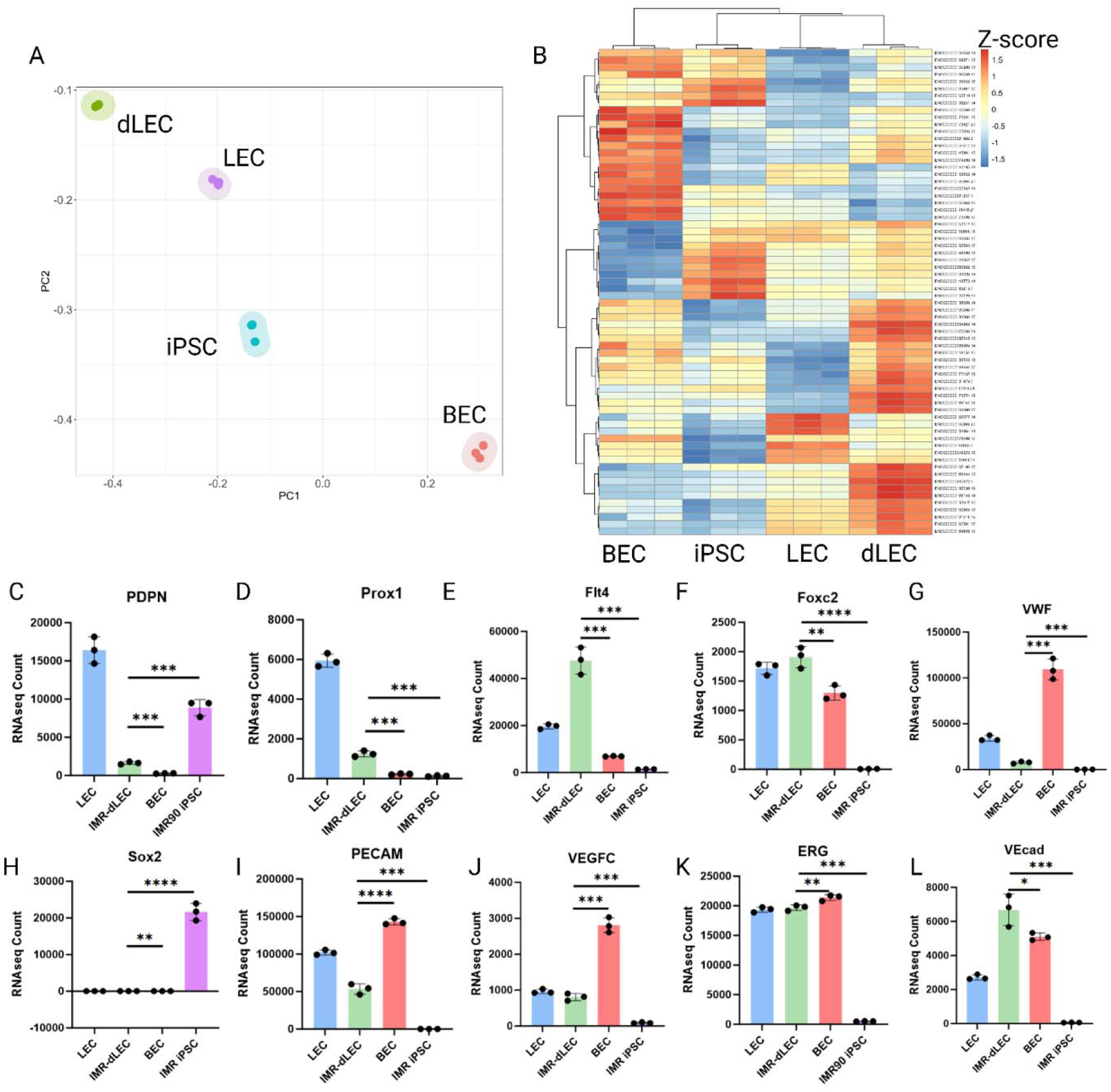
Bulk RNA sequencing results comparing hiPSC, BEC, LEC, and dLEC. (A) Principal component analysis from the gene expression matrix result. Each point represents one biological replicate with RNA collected individually. The PCA was performed on the top 1% of the genes with the greatest standard deviation among the mean values for each cell type. (**B**) Heat map of the top 1% of the genes with the greatest variability, plotted ith Ward.D2 clustering algorithm. (**C-L**) Select relevant endothelial, lymphatic, and pluripotent marker gene expression levels quantified from the gene expression matrix.

We also analyzed the expression levels of known lymphatic, endothelial, and pluripotency markers and found that our dLECs express comparable levels of all expected markers to that of LECs. Our dLECs express comparable levels of VE-cadherin, PECAM, Foxc2, and ERG. The dLECs also express slightly lower, but still comparable, levels of PDPN, VWF, and Prox1, while expressing significantly higher levels of VEGFR3, most likely due to the direct effects of the sodium acetate treatment (**Figure 4C-L**). They also express no pluripotency markers such as Sox2, indicating that they are successfully differentiated into LECs.

Interestingly, hiPSCs have been shown to express PDPN at high levels, which was previously not reported in literature **(Figure 4C)**. However, all fibroblast-derived iPSC cell lines reported by the Human Induced Pluripotent Stem Cell Initiative to the European Bioinformatics Institute show high levels of PDPN expression, indicating that the PDPN expression may be a previously unreported characteristic of fibroblast-derived iPSCs **(Supplementary Figure 10)**. However, this PDPN expression is lost as the cells undergo differentiation, as demonstrated in the differentiated ECs only with VEGF-C and without sodium acetate (**Figure 2G**). Likewise, a differential gene expression dataset comparing iPSCs to iPSC-derived cardiomyocytes show that PDPN expression reduces significantly as the cells differentiate **(Supplementary Figure 11)**.^41^ Therefore, published datasets and our data suggest that PDPN expression may be lost in the differentiation process as hiPSCs exit the pluripotent stage, and reintroduced as the differentiated ECs undergo lymphatic identity induction. As far as the authors are aware, this expression of PDPN in fibroblast-derived iPSC is unreported in literature as of date.

### Differentiated LECs secrete similar cytokines compared to native LECs

In addition to their role in forming lymphatic vessels, LECs have also been known to secrete cytokines that is responsible for the paracrine crosstalk between LECs and the surrounding cells. For example, LECs secrete reelin that promote cardiac organ maturation, and can also secrete CCL21 that promote the growth of glioblastoma stem cells.^11,42^ Pathological LECs, such as LECs exposed to tumors, have also been shown to exhibit abnormal secretion of IL6 which promote primary tumor growth.^43^ These findings suggest that the cytokine secretion is an important aspect of LEC functionality that influences the overall tissue development or tumor growth.

Therefore, we test for the cytokine production of our dLECs and compare them to native LECs and found that all dLECs secrete similar levels of reelin as LECs (**Figure 5A**).^44^ While several cells such as macrophages and fibroblasts have been known to secrete VEGF-C, LECs have not been found to secrete significant amounts of VEGF-C. As such, we found extremely low expression of VEGF-C secretion from all cell lines in the range of picograms per mL, as expected (**Figure 5B**).^45,46^

**Figure 5.**
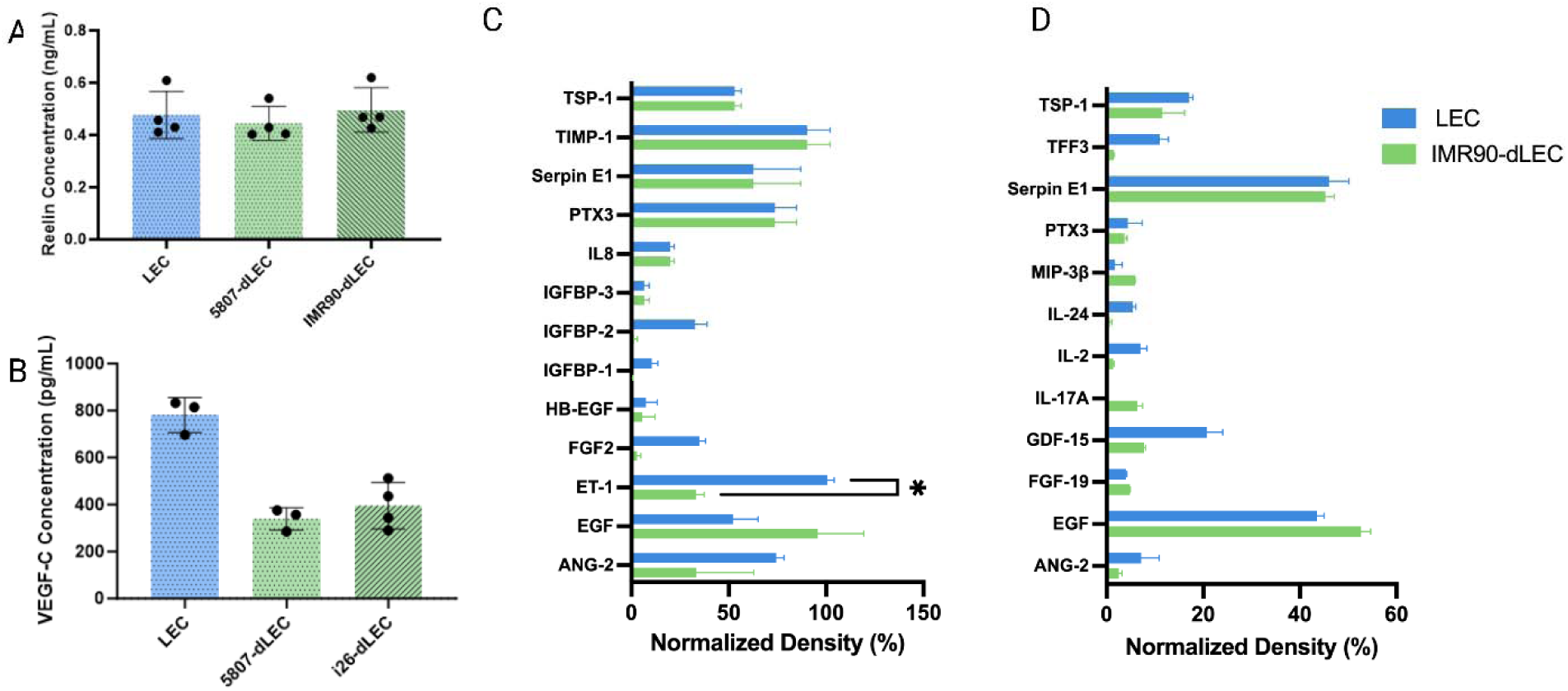
Cytokine and secretomic analysis. (**A**) Reelin ELISA analysis on the supernatants for native and differentiated LECs. ANOVA result showed no significance. (**B**) VEGF-C ELISA analysis on the same supernatant samples. All samples showed secretion levels below 1 ng/mL. (**C-D**) Angiogenesis-related and inflammation-related cytokine array assay results. Only those with measurable level of signal detected via chemiluminescence is shown. Student t-test were performed for each cytokine type, with only ET-1 showing significance. Data represent mean SD, with *n* = 3-4 for ELISA and *n* = 2 for cytokine array.

The proteome profiler assays demonstrated the presence of various lymphangiogenesis-related and anti-inflammatory proteins in both LECs and dLECs that are indicative of their potential therapeutic effect *in vivo* (**Figure 5, Supplementary Figure 12**). The prominent analytes, such as Serpin E1 (PAI-I), has been associated with improving lymphedema.^47^ Pentraxin 3 (PTX3) has been found to be essential for LEC/vessel sprouting as well as the organization and function in lymphatic system.^48^ Epidermal growth factor (EGF) has also been found to promote lymphatic vessel formation.^49^ The presence of other lymphangiogenesis-related factors, such as angiopoietin-2 (Ang-2) and thrombospondin-1 (TSP-1), was also found in both LECs and dLECs thus further demonstrating its potential regenerative functionality *in vivo*.^50–53^

### Differentiated LECs retain network formation capability of native LECs

To demonstrate that our dLECs retain the *in vitro* network formation capability, we compared the networks formed by native and differentiated LECs on two pro-angiogenic natural hydrogels, Matrigel and fibrin gels. Matrigel-based angiogenesis, first described in 1988, is a widely used standard to characterize the angiogenic capacity of endothelial cells, which is crucial for their application in tissue engineering.^54^ Subsequent research has also identified other pro-angiogenic biomaterials that promote *in vitro* network formation and therefore has been used as a tool to measure the angiogenic potential of endothelial cells.^55,56^ LECs have also been shown to form lymphatic networks similarly to their blood counterparts, and even maintain their ability to discriminate between blood and lymphatic lineage cells to form distinct networks when cultured together.^29^

We performed the network formation assays on growth factor reduced Matrigel and fibrin gel substrate, placed on a double-welled ibidi μ-slide. The cells, pre-stained with CellTracker Green CFMDA dye, were allowed to form networks for up to 24 hours, with imaging taking place every hour. Matrigel-seeded cells reached full network formation around 6 hours after seeding, and the fibrin-seeded cells reached its full network around 18 hours after seeding. The representative images of these networks at 6h and 18h are shown for both the LEC and dLEC, showing similar degree of vascularization (**Figure 6A, D**). We quantified the networks using AutoTube, a MATLAB-based software that performs skeletonization of the networks and performs quantification on the identified network.^57,58^ We observed that the dLECs and LECs formed comparable networks, with similar branch point count and vessel width (**Figure 6B-C** and **E-F**). Interestingly, the dLECs formed slightly wider vessels on fibrin, but was comparable on the Matrigel. Overall, these results suggest that our dLECs developed the angiogenic potential that matches that of native LECs.

**Figure 6.**
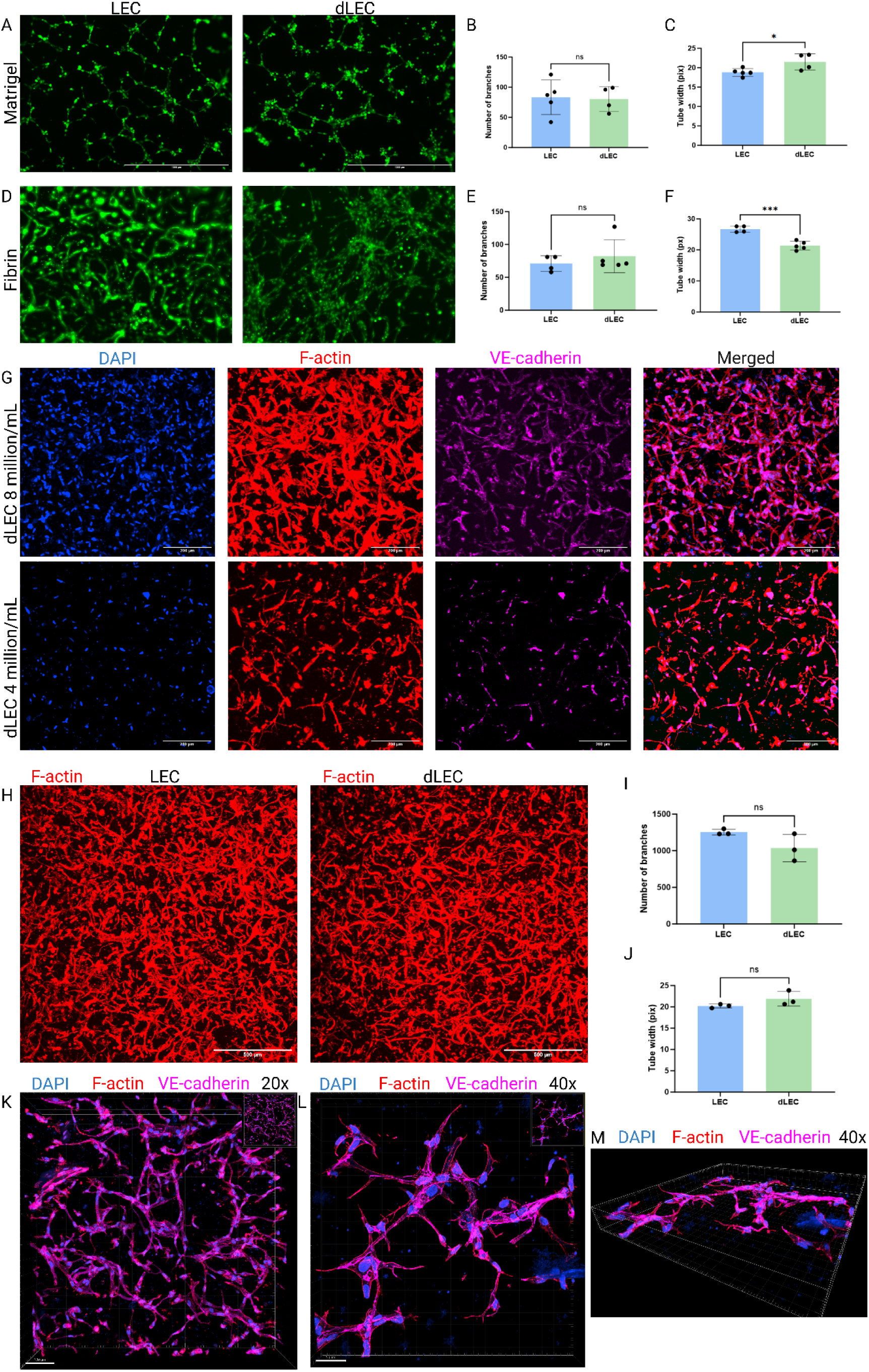
2D and 3D *in vitro* lymphatic vasculature formation. (**A,D**) Representative images of dLECs and native LECs forming networks in Matrigel and fibrin hydrogel surfaces. Selected images were taken 5 hours after initial seeding. Scale bar represents 1000 μm. (**B,C,E,F**) Quantification of the networks for number of branches and network width showing comparable levels between dLEC and LEC. G) Z-max projection images of dLEC vasculature in NorHA seeded at 4 and 8 million cells/mL staining for F-actin and VE-cadherin. Scale bar represents 200 μm. (**H**) Z-max projection of dLEC and LEC vasculature in NorHA stained for F-actin. Scale bar represents 500 μm. (**I-J**) Quantification of the max projection images for network branch count and tube width. (**K-M**) 3D rendering of the dLEC-NorHA vasculature showing VE-cadherin junctions and 3D structures. All statistical analysis performed using Student’s *t*-test. *p<0.05, ***p<0.001.

### Differentiated LECs generate robust 3D lymphatic networks in synthetic hyaluronic acid hydrogels

Despite its widespread adoption, the traditional angiogenesis assay is limited by the 2D nature of the cell growth, which may not accurately capture the angiogenic environment *in vivo.* Therefore, we explored encapsulating the cell suspended in a hydrogel and analyzing the 3-dimensional (3D) network formation that better mimics *in vivo* angiogenesis. Specifically, LECs uniquely express LYVE-1, a lymphatic-specific homologue of CD44, which binds to hyaluronic acid to mediate lymphangiogenesis.^59,60^ Our previous works have shown that hyaluronic acid hydrogels not only promote lymphatic vessel formation, but also is capable of modulating the lymphangiogenesis through stiffness modulation.^58,61^

For this study, we utilized hyaluronic acid hydrogels modified with norbornene (NorHA) which enabled UV-inducible thiol-norbornene crosslinking, along with matrix metalloproteinase (MMP)-sensitive crosslinker peptides to allow for matrix degradation and subsequently lymphatic network formation.^61,62^ The cells were suspended into hyaluronic acid solution at a concentration of 4 and 8 million cells/mL, which were the optimal density range identified in our previous published work.^63^ Following the UV-mediated crosslinking, the gels were cultured in media supplemented with 100 ng/mL of VEGF-C and FGF-2. We observed an extensive network formation by day 3 after crosslinking with dLEC-encapsulated gels, at which point the gels were fixed and stained for DAPI, F-actin, and VE-CAD (**Figure 6G**). As expected, the 8 million/mL gels formed more robust vascular network due to the higher cell density, while the 4 million/mL gels demonstrated a sparser network. We repeated the gel formation with native LEC and dLEC and stained for F-actin and found similar degree of network formation between dLEC and LEC (**Figure 6H-J**).

The 3D rendering of these vessels show that these vessels are indeed 3D and extend across the entire volume of the gel (**Figure 6K**). At 40x magnification, we see additionally that the dLECs are forming functional junctions based on the staining expression of VE-cadherin around the edges of the cell, along with the F-actin expression along the vessel formation (**Figure 6L**). We can additionally see that the vessels form a three-dimensional structure that closely mimic lumen and lymphatic button-like junctions, indicating that our dLECs are capable of forming highly accurate and robust lymphatic vasculature *in vitro* (**Figure 6M, Supplementary Movie S1**). These results suggest that our dLECs, similar to native LECs, can form robust lymphatic vasculature in an *in vitro* setting, which will allow them to be used in forming engineered tissues with functional lymphatic vessels.

### Differentiated LECs promote lymphatic regeneration in mouse models

One of the potential applications of dLECs is in developing therapeutic options for secondary lymphedema. Currently, lymphedema treatments are limited to compression, anti-inflammatory drugs, and lifestyle changes.^64–66^ Recent developments in lymphedema therapeutics include various pro-lymphangiogenic growth factors, indicating that promoting lymphangiogenesis may be a viable method to restore lymphatic functionality in secondary lymphedema.^67–72^ Therefore, we explored the use of dLECs as a novel and fully autologous method to promote lymphatic regeneration in two relevant murine models: double-ligation tail lymphedema model and mammary fat pad model.

In the first murine model of double-ligation tail lymphedema model, we induced artificial lymphedema in mouse tails by ligating the two collecting lymphatic vessels and all surface lymphatics. This tail lymphedema model has been widely used and adapted for modeling secondary lymphedema by disrupting the lymphatic function across the tail, thereby causing swelling in the distal side of the tail similar to the symptoms of secondary lymphedema.^73,74^ The disruption of the collecting vessels was shown to be sufficient to induce drastic lymphatic remodeling, with swelling taking up to 4 weeks to return to pre-injury levels.^75^ In this study, we performed similar ligation of both collecting vessels and all superficial lymphatics and verified that the swelling reaches maximum level in about a week, which can be prevented through daily injection with non-steroidal anti-inflammatory drugs (NSAID; **Supplemental Figure 13**).

We injected dLECs suspended in Matrigel at concentration of 10 million cells/mL directly at the wound site and allowed the Matrigel to solidify for up to 15 minutes on a heating pad. The treatment group was compared with mice with no injection and mice with only Matrigel injected without cells. NSAID-treatment group was included as a positive control. We sacrificed the mice at day 7 to evaluate the efficacy of the dLECs in reducing swelling and restoring lymphatic vessels (**Figure 7A**).^76^ By day 5, a significant difference in swelling between the treatment and control groups was observed (**Figure 7B-D**). The mice that received Matrigel-dLECs injection experienced around 20% of the pre-injury level which was statistically non-significant compared to the NSAID-treatment group (**Figure 7E, Supplemental Figures 14-17**). In comparison, the mice with blank Matrigel injection and no injection experienced similar degree of swelling of around 40% increase from the pre-injury levels. Interestingly, the mice that received cell injection experience a sudden increase in swelling around day 7, which suggests initial immune reaction to xenogenic cells.

**Figure 7.**
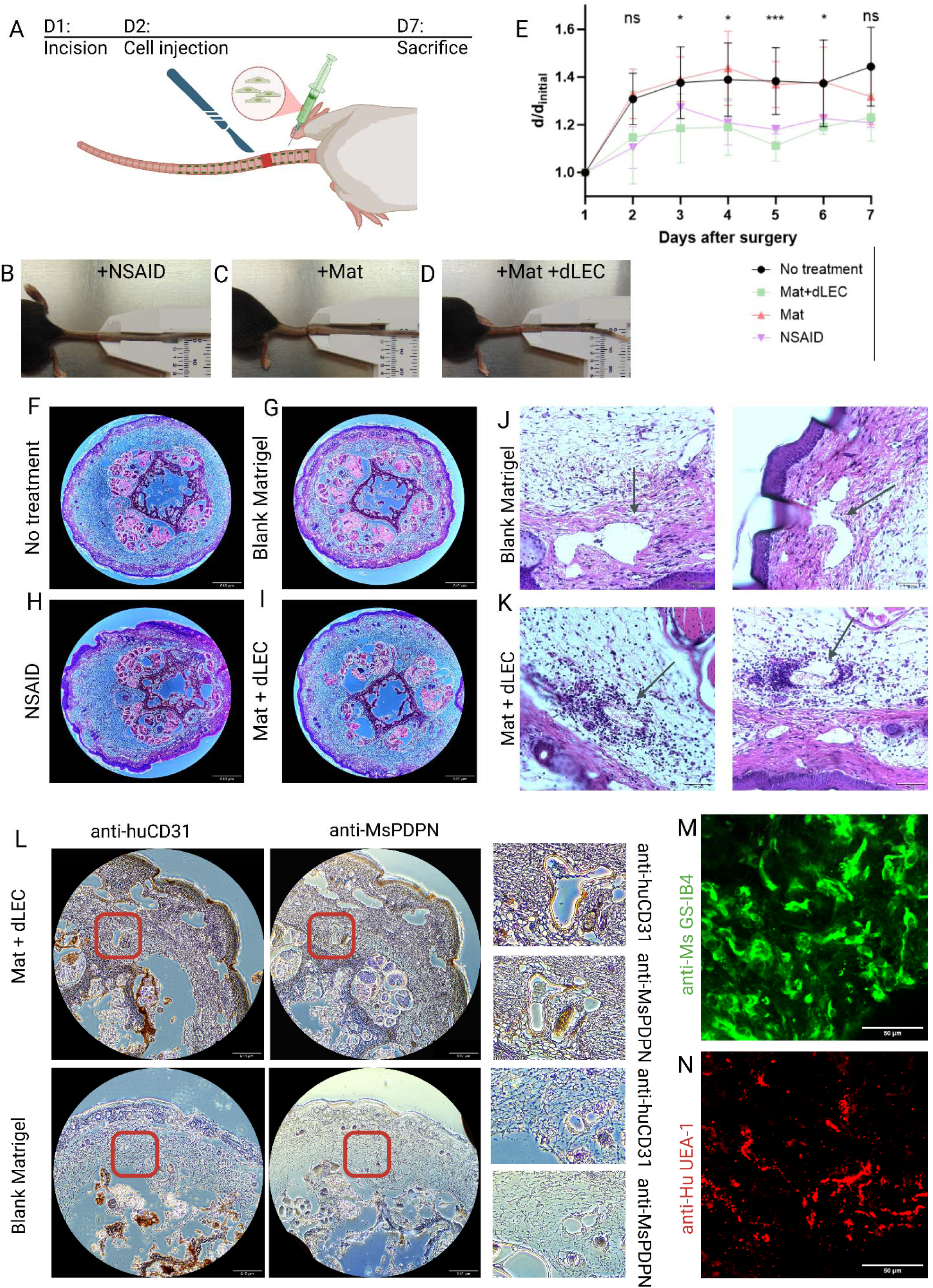
*In vivo* functionality of differentiated LECs in murine models. (**A**) Schematic of the mouse tail lymphedema model. (**B-D**) Representative images of mice at day 5 following the surgery showing the differences in the degree of inflammation. (**E**) Changes in tail diameter for the 7 days following the surgery, normalized to the initial diameter. All statistical analysis is performed between the treatment dLEC group and blank Matrigel group. (**F-I**) H&E stains of the cross-sections of the wound site showing full disruption of the collecting lymphatics and most surface lymphatics. Scale bar is 860 μm. (**J-K**) Higher magnification images of the H&E staining focusing on the lymphatic lumens showing the presence of the dLECs in the wound site. Scale bar is 70 μm. (**L**) Immunohistochemistry slides of cross sections of the tail proximal to the wound site for blank Matrigel and Matrigel-dLEC treated group stained for mouse PDPN and human CD31, with areas of interest zoomed in. Positive stain is shown in brown. Scale bar is 860 μm. (**M-N**) Z-max projection images of the granular hydrogels stained with rhodamine-conjugated *Ulex Europaeus Agglutinin I* (*UEA-1 lectin*) for human endothelial cells and fluorescein-conjugated *Isolectin Griffonia simplicifolia* (*GS-IB 4* isolectin) for mouse endothelial cells. Scale bar represents 50 μm. All statistical analysis performed using Student’s *t*-test. *p<0.05, ***p<0.001.

We sacrificed the mice at day 7 and collected the wound site sample from each mouse. We prepared histology samples sliced along the direction of the tail at the distal side of the wound and performed H&E staining. In all samples, we observe no major structures that indicate collecting lymphatic vessels, indicating that we have successfully disrupted the collecting vessels (**Figure 7F-I**). Since our dLECs mimic dermal LECs and do not contain necessary components of a collecting vessel such as lymphatic valve and smooth muscle cells, we do not expect the regeneration of lymphatic collecting vessels within this timeframe. We identified several lumen-like structures that are most likely superficial dermal lymphatics based on the presence of endothelial barrier along the edge of the lumen and lack of red blood cells to distinguish it from the blood vessels. Within these lymphatic-like lumens, we notice a higher deposition of elongated endothelial-like cells and macrophage-like multi-nucleated cells, indicating our dLECs are present within the lymphatic lumens (**Figure 7J-K**).

To further confirm the presence of our dLECs in the wound site, we collected histology samples distal to the wound site, where we surface lymphatic repair to begin. We performed immunohistochemistry staining for human CD31 and mouse PDPN to identify the interface between human dLECs and mouse LECs. Two representative tail samples, one from the group that received dLECs and one treated with blank Matrigel, are shown stained for mouse-PDPN and human-CD31, with areas of interest emphasized (**Figure 7L, Supplemental Figure 20**). The positive stains are visible around the endothelium of the vessel lumen in both the human-CD31 and mouse-PDPN stains, indicating that the dLECs are contributing to the lymphatic repair process. We confirmed that the mice samples that received a blank Matrigel treatment exhibited few mouse-PDPN-positive vessels but lacked any human-CD31 expression in the vessels, along with the standard IgG negative control stains (**Supplemental Figures 18-19**). Lymphatic vessels were identified by the presence of PDPN-expressing endothelial cells surrounding the lumens of interest. In several of the identified lymphatic vessels, we observed chimeric vessels made of human and mouse cells, indicating that our dLECs are able to integrate into the host murine lymphatic vasculature.

To further investigate how human dLECs can integrate with the host murine lymphatic vessels, we used a second mouse model of murine mammary gland model, which has been previously used in the context of patient-derived tumor xenograft models of breast cancers.^77^ Granular hydrogel constructs were fabricated using NorHA and cultured *in vitro* for two days to allow dLECs to begin to elongate and sprout, and then they were implanted into a pocket created in the #4 mammary gland of NOD/SCID mice.^61,78^ To assess lymphatic integration, as well as to distinguish mouse vasculature from the implanted human LEC networks, two lectin dyes to label endothelial cells (ECs), one specific to mouse and the other one specific to human ECs, were injected into the extracellular space of the hindfoot for uptake to the lymphatic circulatory system before tissue harvest. The fluorescein-conjugated *Isolectin Griffonia simplicifolia* (*GS-IB_4_* isolectin) stained mouse ECs, both blood and lymphatic, and the rhodamine-conjugated *Ulex Europaeus Agglutinin I (UEA-1* lectin) stained the human LECs^42,43^. While *UEA-1* lectin can stain both blood and lymphatic human ECs, the entirety of the rhodamine signal in these explants can be attributed to the implanted human LECs due to using a monoculture hydrogel system. In the granular gels, we observe vascular-like structures forming similar to the *in vitro* results we have published previously, which integrates with the mouse endothelial cells (**Figure 7M-N**). Taken together, these results suggest a potential therapeutic option for secondary lymphedema using patient-derived dLECs.

## Discussion

Since the discovery of lymphatic-specific markers that made lymphatic vessel isolation possible, there has been an ever-increasing focus around the lymphatic system.^16,79^ Despite its relative obscurity compared to its well-known blood circulatory counterpart, lymphatic vessels have been shown to play an indispensable role in critical tissue functions, such as fluid homeostasis and immune cell trafficking.^1–3^ Therefore, lymphatic endothelial cells, which line the lymphatic vessels distinct from blood vessels, have been subject of focus for tissue engineering researchers with the intent to incorporate the lymphatic system into engineered tissue constructs. With the rise of stem cell engineering that aims to develop new alternative methods (NAMs) for personalized therapies and drug screening platforms, the demand for hiPSC-derived LECs has grown, but to date, there is a lack of xeno-free and efficient differentiation method that captures both the lymphatic marker expression and LEC functionality.

We previously reported a direct differentiation of hiPSCs into LECs using transcription factor *ETV2* and *ETS2*.^80^ While direct differentiation protocol can be used for disease modeling, the lentivirus used in the protocol may not be suitable for clinical applications. To complement this direct differentiation protocol, here we reported a step-wise differentiation using metabolic pathway inspired by the existing hiPSC-EC differentiation protocol, which utilizes a sequential differentiation from hiPSC to endothelial progenitor-like cells, then to mature endothelial cells.^36^ Interestingly, we have found that an indirect differentiation method from endothelial progenitor-like cells to lymphatic endothelial cells yielded significantly higher lymphatic population compared to a direct lymphatic differentiation from mesodermal state. This is potentially due to the protocol closely following the centrifugal theory of lymphatic origin, which posits that the lymphatic vessels develop out of blood vessels during embryogenesis.^81^

We have shown that with combination of VEGF-C and sodium acetate treatment during the lymphatic induction stage, there is a sufficient upregulation of VEGFR3, which then upregulates Prox1 and subsequently further downstream lymphatic markers such as PDPN. We also showed that our cells retain the generic endothelial markers such as CD31, VE-Cad, ERG, VWF, and others. Though our iPSCs unexpectedly expressed high level of PDPN, published datasets suggest that this is characteristic of fibroblast-derived iPSCs and PDPN expression is subsequently lost as iPSCs exit their pluripotency state until the cells mature into lymphatic identity. When compared to the currently widely accepted differentiation protocol using the OP9 cell line, our differentiation protocol results in higher lymphatic marker expressions while also exclusively relying on xeno-free and fully defined components which improve the clinical applicability of these dLECs.

In addition, we have shown that our dLECs retain most of the functionality of native LECs, including cytokine production, network formation, and lymphangiogenesis. In order to successfully integrate into engineered tissues, LECs are required to form lymphatic vessels which can then fulfill the native lymphatic functions such as fluid transport. Here, we showed that lymphangiogenic capacity of our dLECs match that of native LECs, in both traditional 2D angiogenesis assays and the novel 3D vessel formation assay that requires complex functions such as junction formation and matrix degradation. Furthermore, we show that the dLECs also have a similar cytokine secretion profile, which is crucial for surrounding organ development in *in vitro* organogenesis. We have shown that our dLECs are equally functional to native LECs, which demonstrates the potential for the dLECs to be incorporated into engineered tissues in order to model the lymphatic system.

We also demonstrated the potential therapeutic value of the dLECs through the mouse tail lymphedema model. Injection of the dLECs into mouse tail accelerated the lymphatic vessel repair, indicating that the dLECs are able to incorporate into the reconstructed lymphatic system. These results suggest that the dLECs, in addition to their value in creating *in vitro* lymphatic vessels, can also have direct therapeutic applications in lymphatic diseases such as lymphedema, where current treatment options are limited and primitive.^82^ Taken together, we conclude that FAO-metabolite supplementation drives lymphatic identity upregulation in differentiated ECs that approach the native LECs in functionality and characteristics. Future works should explore the use of dLECs in applications such as organoid models and organ-on-a-chip models to incorporate the indispensable functions of the lymphatic system into tissue engineering,^83,84^ as well as to evaluate the efficacy of these dLECs in clinically-relevant models of lymphedema.^85^

### Limitations of the Study

Recent discoveries suggest that there are two distinct subpopulation of LECs, one arising from endothelial progenitor cells and one differentiating directly from mesenchymal stem cells.^30^ The endothelial progenitor-derived LECs correspond to the traditional centrifugal theory where LECs are thought to be developed from early blood vasculature during embryogenesis, while the mesenchymal-derived LECs correspond to the centripetal theory where lymphatic vessels are thought to develop independently in an organ-specific manner.^81,86^ This dual origin of LECs means that LECs are heterogenous in nature, with certain organ-specific LECs such as meningeal or lacteal lymphatics derived from local mesenchymal stem cells expressing organ-specific markers and functions.^87,88^ In our differentiation protocol, we closely follow the centrifugal theory, which generates LECs with generic lymphatic markers similar to endothelial progenitor-derived LECs, but lack certain organ-specific lymphatic subtype features.

While our differentiation protocol is inspired by the lymphogenic process during embryogenesis, practical constraints force us to greatly expedite this differentiation process *in vitro*. In human embryogenesis, lymphatic vessels take up to 30 days to first express Prox1, and subsequent maturation is a years-long process that lasts through birth and childhood.^89^ Furthermore, lymphatic vasculature requires the mechanical stimulation from the environment, such as pressure or shear stress, to undergo maturation into lymphatic valves.^90,91^ Therefore, the differentiated LECs that we generate are most likely immature compared to native LECs collected from post-natal donors. Future studies should explore maturation of dLECs through combination of biochemical and mechanical cues.

Due to cost constraints, there is a limit to the number of hiPSC cell lines that is available at our disposal. Therefore, this protocol should be repeated using various hiPSC cell lines that have been derived from a variety of sources and reprogramming methods to account for potential effects of epigenetic memory.^92^ The *in vivo* study is limited by the lack of mice strain variants, and subsequent experiments investigating the restoration of lymphatic fluid transport in larger animal models are likely required to fully ascertain the therapeutic potential of the dLECs.^85^

## Methods

### Human iPSC and LEC Culture

Human induced pluripotent stem cells derived from either skin or lung fibroblasts or peripheral blood mononuclear cells (WiCell, Stem Cell Technologies, ATCC) were expanded and used for experiments for up to 15 passages from initial purchase (**Supplementary Table 1**). Human iPSCs were seeded on tissue culture plastic surfaces coated with 0.083 mg/mL of Matrigel (Corning) in DMEM-F12 media (Gibco) and grown in mTeSR+ media (Stem Cell Technologies) at 37°C with 5% CO_2_. Between 2 to 10 μM of Y-27632 Rho-kinase inhibitor was added to the cell culture media for the first day after seeding depending on the iPSC cell line requirements, then subsequently removed. Human LECs and BECs derived from the dermis of two adult donors (Promocell) were expanded and used between passages 5 and 8. The cells were cultured in MV2 media (Promocell) and similarly incubated at 37°C with 5% CO_2_. LECs were characterized for the positive expression of CD31 and Prox1 throughout the experiments. All cell lines were routinely tested for mycoplasma contamination and were negative throughout this study.

### Human iPSC Differentiation

Human iPSCs were passaged at between 60-80% confluency using Accutase and separated out into single cell suspension. The cells were passaged onto a new Matrigel-coated plate at a concentration of between 30,000 and 50,000 cells per cm^2^ of cell growth area. The exact seeding density was optimized for each cell line. Cells were treated with 2-10 μM of Y-27632 Rho-kinase inhibitor (Stem Cell Technologies) in mTeSR+ media (Stem Cell Technologies) for 1 day after passaging. Following this, cells were switched to APEL2 media (Stem Cell Technologies) containing 6 μM of CHIR99021 GSK inhibitor (Stem Cell Technologies) for 2 days without media change. On day 3, cells were switched to APEL2 media containing 50 ng/mL of VEGF-A (Stem Cell Technologies), 25 ng/mL of BMP-4 (R&D Biosystems), and 10 ng/mL of FGF-2 (R&D Biosystems) for 2 days, with media change each day.

At day 5, cells were passaged using TripLE Express media and re-seeded onto tissue culture plastic at density of 60,000 to 100,000 cells per cm^2^. The cells were cultured in MV2 media (Promocell) containing 100 ng/mL of VEGF-C (R&D Biosystems) and 50 mM of sodium acetate (Thermo Fisher). Cells were cultured for 4 days with media change after 2 days. Following the differentiation, cells were collected for further experiments or used for RNA isolation.

### Gene Expression

To analyze the gene expressions, we collected the RNA from dLECs on day 9 of differentiation, or 4 days after seeding for LEC and BEC. Each sample contained at least 3 biological replicates from cells cultured in distinct wells and analyzed with qRT-PCR. RNA was isolated using Qiagen Mini-prep kit (Qiagen), and quantified and diluted to the same concentration for all samples. The RNA was reverse transcribed using High-capacity cDNA reverse transcription kit (Thermo Fisher) using the recommended protocol from the manufacturer. We used TaqMan Gene expression assays (Thermo Fisher) following manufacturer protocols and used *GAPDH* and *beta-actin* as endogenous controls to normalize relative gene expression levels using the ΔΔC_T_ method.

### Flow Cytometry

For flow cytometry analysis, cells were detached using TripLE Express as described in previous sections, then subsequently fixed using 4% paraformaldehyde solution for 15 minutes in 4°C. Then the cells were washed and stained in eBioscience FACS buffer (Thermo Fisher) containing the recommended amount of conjugated antibody solution. For nuclear marker staining such as ERG and Prox1, eBioscience Intracellular Fixation and Permeabilization buffer set (Thermo Fisher) was used instead. For each experiment, unstained control was included as a negative control. The samples were run using FACSMelody Cell Sorter (BD Biosciences) or Cytek Northern Lights CLC (Cytek) until at least 5,000 events were captured. The list of antibodies used is listed in Supplemental Table 3.

The resulting data was processed by removing doublets using FSC-A vs FSC-H plot and removing the smaller cluster above the linear main population corresponding to non-single cell population. The fragments were removed using FSC-A vs SSC-A plot, and removing the events expressing very low FSC or SSC. For gating the populations, unstained controls for each stain were used to identify the positive-negative expression boundaries. Identical gates were used for each replicate and sample for each individual run. Each sample except unstained controls was run in biological triplicates where cells were cultured and differentiated separately, then stained separately.

### 2D Network Formation Assay

The LECs and dLECs were first cultured to 70-80% confluency, then stained with 5μM CellTracker Green CFMDA (Thermo Fisher) for 30 minutes. The cells were passaged 24 hours later and seeded at a density of 30,000 cells per cm^2^ onto the Ibidi 15 well μ-slide (Ibidi) containing either Matrigel or fibrin gel. The Matrigel was prepared by placing 10μL of growth factor reduced Matrigel (Corning) thawed out on ice to 4°C, then incubating at 37°C for 30 minutes. The fibrin gel was prepared by mixing 6μL of the fibrinogen solution with the 4μL of thrombin solution from the Angiogenesis Assay Kit (Millipore Sigma), and incubating at 37°C for 30 minutes. The cells were allowed to settle and attach for 1 hour after seeding, then imaged every hour for up to 24 hours.

### Network Quantification

The fluorescent channel images from each well in the μ-slide were pre-processed by performing a rolling ball background subtraction with radius of 50 pixels on ImageJ. Then the resulting images were used as inputs to the AutoTube software and skeletonized. The skeletonized images were manually checked, and corrected as needed to remove holes in networks due to thresholding error. The resulting skeletonized images were analyzed using default settings, and the branch point and tube width data were collected for each replicate in each group.

### RNA Sequencing Analysis

RNA samples were prepared using the Qiagen Mini-prep kit and further purified as needed to reach minimum A260/230 ratio of 1.8. The RNA samples were subsequently used for library prep using Illumina NextSeq P1 600 cycle at minimum read depth of 50 million per sample. The resulting sequencing readouts were aligned with the human genome using the STAR RNA-seq aligner, and the gene count matrix was generated.^93^ The gene count matrix was normalized, then the top 1% of genes with the highest differential expression were selected. The resulting data matrix was used to perform the principal component analysis and generate a heat map, where the genes with the greatest differential expression between the control and the experimental groups were selected. The gene counts for several key genes of interest were extracted and subsequent statistical analyses were performed.

### Cytokine ELISA

Supernatants were collected and filtered from confluent cultures of either LEC or dLEC. Media was changed 24 hours prior to supernatant collection to MV2 growth media (Promocell) that does not contain any additional growth factors. Supernatants were centrifuged at 300g for 5 minutes to pellet any cell debris. We quantified the relative total protein concentration using Bradford Protein Assay (Thermo Fisher) and diluted samples to the same concentration using PBS. The normalized supernatants were then quantified for cytokine content using ELISA kits for human VEGF-C and human reelin (Abcam). The absorbance values were quantified using Spark Microplate Reader (Tecan).

### Proteome Profiler Arrays

Supernatants were collected and filtered from confluent cultures of either LEC or dLEC. Media was changed 24 hours prior to supernatant collection to MV2 growth media (Promocell) that does not contain any additional growth factors. Supernatants were centrifuged at 300g for 5 minutes to pellet any cell debris. The Proteome Profiler Human Angiogenesis Array Kit (R&D Systems) and Human XL Cytokine Array Kit (R&D Systems) were used to detect the expression of multiple proteins. Briefly, the supernatants were mixed with corresponding cocktails of biotinylated detection antibodies and incubated for 1 hour at room temperature. The mixtures were then incubated with array membranes containing specific capture antibodies to kit-specific target proteins overnight at 4°C on a plate shaker. The membranes were then washed to remove unbound material and incubated with streptavidin conjugated to horseradish-peroxidase (HRP) for 30 minutes at room temperature on a plate shaker. The membranes were then washed to remove excess streptavidin-HRP and were prepped with chemiluminescent detection reagents to be imaged using a ChemiDoc-It2 imager (UVP, Analytik Jena) and VisionWorks software (Analytik Jena). Quantification of the integrated pixel density of the duplicate spots was done using ImageJ and the MicroArray Profile plugin (OptiNav Software). Values from each membrane were normalized to an average pixel density value from respective reference spots.

### Norbornene-modified Hyaluronic Acid Hydrogel Formation

Norbornene-modified hyaluronic acid polymers were synthesized using methods described in our previous publication with a 19% substitution of available functional groups with norbornene (19% D.S.).^62^ The polymer was weighed out and mixed with PBS to the concentration of 2 wt%. Additionally, 0.5mg/mL of Irgacure 2959 (Sigma-Aldrich), 5 mM of RGD peptide (Genscript) and 2.6 mM of MMP-sensitive crosslinker (Genscript) were added to the mixture. The cells were passaged and resuspended in the polymer mixture. The polymer solution was placed into a mold of 50μL in volume and placed under 254 nm UV light at 10 mW/cm^2^ intensity for 70 seconds to induce crosslinking, then placed into MV2 media (Promocell) containing 100 ng/mL of recombinant human VEGF-C (Bio-Techne), 100 ng/mL of recombinant human bFGF (Stem Cell Technologies), and Antibiotic-Antimycotic (Thermo Fisher). Media was changed every day without disturbing the gel.

### Immunofluorescence assay of hydrogels

Hydrogel samples were carefully placed on glass-bottom dish and fixed in 4% paraformaldehyde solution for 30 minutes at room temperature on the shaker. The gels were then blocked with 3% BSA solution for 1 hour at room temperature or overnight at 4°C on the shaker. Gels were stained with 300 nM DAPI solution for 30 minutes, then stained with 1:1000 dilution of the phalloidin-AF594 (Thermo Fisher) and VE Cadherin antibody conjugated with AF488 (Santa Cruz Biotechnology) on the shaker overnight. Between each stain the gels were washed 5 times with PBS, each with 5 minutes on the shaker. The gels were imaged on the Nikon AxR Confocal Microscope at either 10, 20, or 40x objective lenses. The resulting fluorescent images were used to generate 3D renderings using the IMARIS software.

### Mouse tail lymphedema model

All animal studies were conducted in accordance with the relevant federal and local regulations and University of Notre Dame’s IACUC protocol (protocol number 24-07-8678). C57BL/6J mice (The Jackson Laboratory) were anesthetized and the tail area sterilized using isopropanol and chlorohexidine wipes. The initial incision was made approximately 15mm away from the base of the tail around the entire circumference, while taking care to not sever the veins underneath the lymphatic collecting vessels. Any mice with noticeably heavy bleeding or necrotic tails, which indicate damage to the veins, were excluded from the study. Following the incision, the entire wound area was cauterized using a Bovie high temperature cauterizer (Aspen Surgical), then a topical antibiotic ointment was applied to the wound. Mice were treated with up to 0.05mL per 20 g weight of Ethiqa XR (MWI Animal Health), and the NSAID groups were treated with up to 0.25mg per 20 g weight of meloxicam (MWI Animal Health). NSAID was administered every 24 hours and Ethiqa XR was administered as needed 72 hours after the initial surgery.

### Mouse tail Matrigel and cell injection

For the Matrigel and Matrigel with cell treatment groups, we injected the respective materials 24 hours following the initial surgery. Growth factor reduced Matrigel (Corning) was thawed out on ice and mixed with dLECs at concentration of 10 million cells/mL. The cell-gel mixture or blank Matrigel were loaded into a syringe with 25-gauge needle and dispensed in the wounded area. The mice were placed on a heating pad under anesthesia for up to 15 minutes to allow for the gel to solidify. The wound area was then wrapped with bandage temporarily to prevent the gels from being disturbed.

### Mouse tail imaging and sample collection

The mice were placed under anesthesia each day and photographed to track the tail swelling diameter. The mice tails were measured using a manual caliper and photographed, while any mice exhibiting necrosis were euthanized and excluded from the study. Following 7 days, the mice were sacrificed using CO_2_ euthanasia chamber followed by cervical dislocation. The section of the mouse tail roughly 20mm long centered around the wound site were collected and placed in 4% formaldehyde solution overnight. The mice tail samples were cryo-sectioned and the H&E stains were prepared.

### Mammary Fat Pad Model

Female NOD/SCID mice (3-4 weeks old) were used for the study as previously described in our previous studies. Briefly, granular hydrogel constructs were maintained in a sterile environment and transferred to the animal facility for implantation into the mice. The granular hydrogels were generated using pipetting emulsification method which was previously published by the authors.^78^ Under isoflurane anesthesia, an incision was made to access both the left and right #4 mammary gland. A pocket was then created with tweezers and the hydrogel construct was inserted into each pocket. Incisions were closed up with staples and mice were kept on a heating pad to recover from anesthesia, before being transferred back to their cage for daily monitoring (*n* = 4 per group). At the end of each experiment, mice were euthanatized using carbon dioxide chamber. We have complied with all relevant ethical regulation of animal use.

Two weeks post-implantation, two lectin dyes to label endothelial cells (ECs) of each species were injected into the extracellular space of the hindfoot for uptake to the lymphatic circulatory system before tissue harvest. The fluorescein-conjugated *Isolectin Griffonia simplicifolia* (*GS-IB_4_*isolectin) stained mouse ECs and the rhodamine-conjugated *Ulex Europaeus Agglutinin I (UEA-1* lectin) stained the human LECs. Hydrogels and the surrounding mammary gland tissue were then harvested. Tissues and hydrogels were arranged in cassettes and fixed with 4% PFA overnight, and then stored in sterile PBS. Due to low signal intensity and challenges with auto-fluorescence from the surrounding adipose tissue, some explants were stained a second time with the lectin dyes after fixation.

Tissues and hydrogels were then imaged as a whole-mount sample in a coverslip chamber-well with a Nikon AXR confocal microscope. The entire tissue area was scanned at 10x to determine the regions of vessel integration, and then z-stacks at 20x were taken to visualize the 3D structure and spatial arrangement of the vasculature. Regions of interest included where the mouse vasculature invaded the hydrogel region and where the human lymphatic vessels within the hydrogel continued to grow and connect with the host vasculature. Regions of fluorescent signal overlap were deemed areas of host-implant integration.

### Statistical analysis

Statistical analysis was performed using GraphPad Prism software (GraphPad Software Inc., La Jolla, CA). All data where statistical analysis is reported were performed with minimum of three biological replicates. Statistical comparisons were made using Student’s *t*-test to analyze the differences between the control and the experimental group. For the multiple comparisons, analysis of variance (ANOVA) was used followed by Tukey post hoc analysis. All results are expressed as mean ± SD, with individual data points shown alongside the mean. Significance levels are reported as *p<0.05, **p<0.01, ***p<0.001, ****p<0.0001.

## Acknowledgements

We acknowledge support from the University of Notre Dame through “Advancing Our Vision” Initiative in Stem Cell Research, Harper Cancer Research Institute – American Cancer Society Institutional Research Grant (IRG-17-182-04), American Heart Association through Career Development Award (19-CDA-34630012 to D.H.-P.), National Science Foundation (2047903 and 2225601 to D.H-P.), National Institutes of Health (1R35-GM-143055 to D.H.-P.), National Science Foundation-Graduate Fellowships Program (NSF-GRFP to D.P.J.). We would like to thank Drs. Sarah Cole and Sarah Chapman from the Notre Dame Integrated Imaging Facility for performing confocal imaging and immunostaining analysis.

## Author contributions

D.P.J. and D.H.-P. conceived the ideas, designed the experiments, interpreted the data, and wrote the manuscript. D.P.J., S.S., D.M., A.T., N.K.L., R.S.G., B.S. conducted the experiments and analyzed the data. J.B.D. helped and consulted with the animal studies. J.B.D. and D.H.-P. supervised the study. All authors have approved the manuscript.

## Data and materials availability

Additional material and data which contributed to this study are present in the **Supplementary Information** and source data can be found in **Supplementary Data 1**.

