## Supplemental Information for "Fatty acid metabolite promotes lymphatic identity in stem cell-derived endothelial cells"

**This PDF file includes:**

Supplementary Figures S1 to S16

Supplementary Tables S1 to S3

**Other Supplementary Materials for this manuscript include the following:**

Movies S1

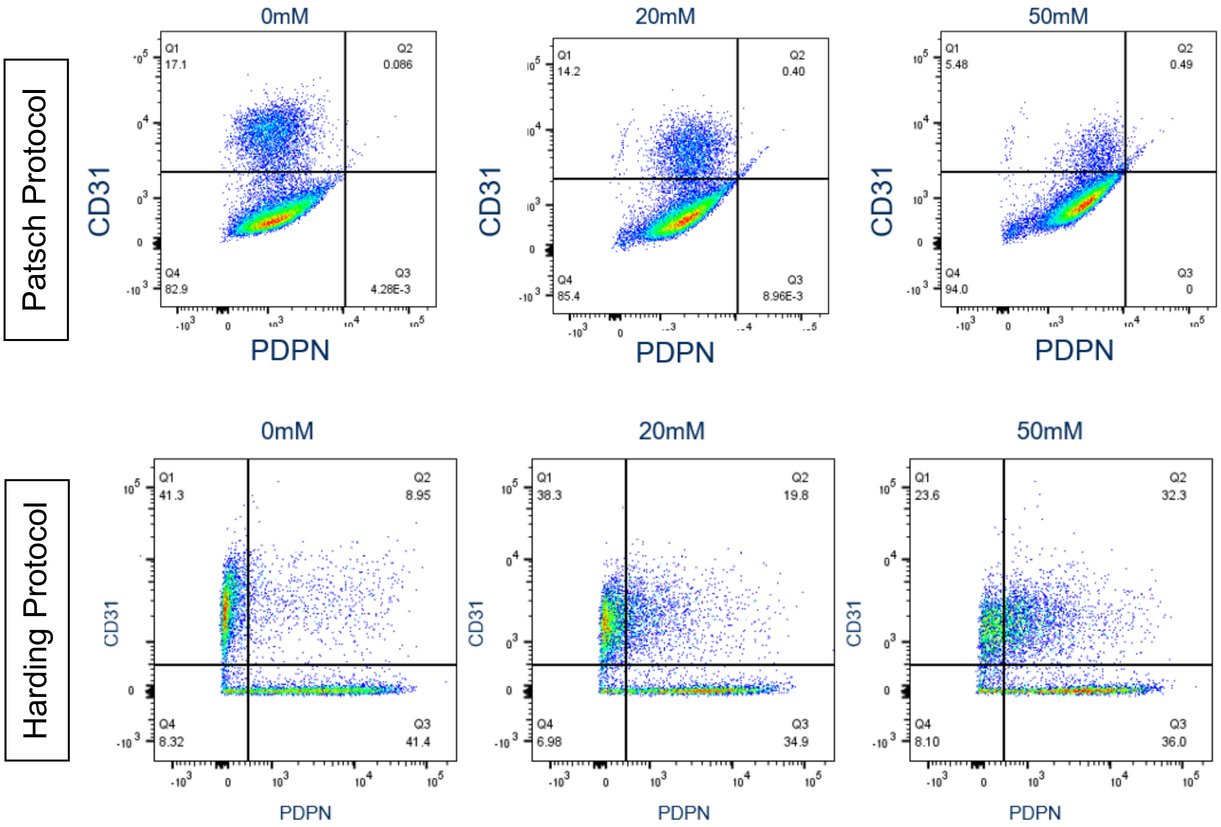


**Supplementary Figure 1**. Comparison between the Patsch and Harding protocols modified with VEGF-C and sodium acetate treatment.

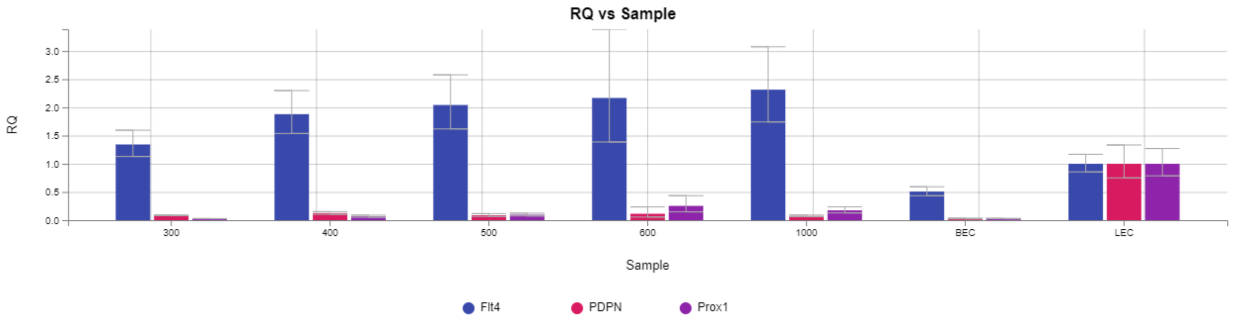


**Supplementary Figure 2**. Relative lymphatic marker expressions for various seeding densities after day 4. The seeding densities are represented as $10^2$ cells/cm$^2$. Three lymphatic markers are shown normalized to P6 LEC expression levels.

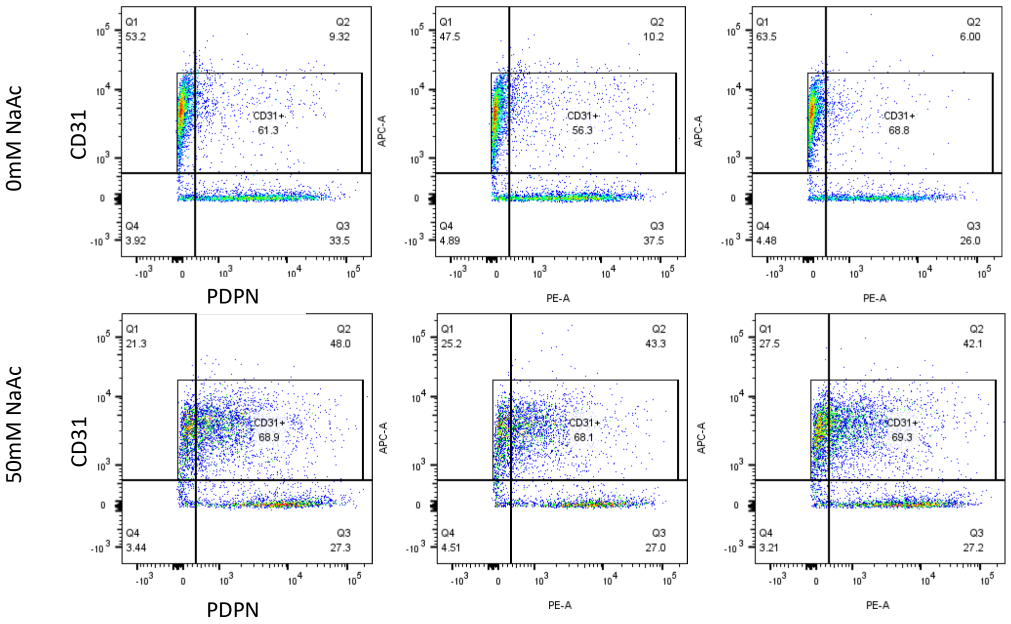


**Supplementary Figure 3**. Human i29 iPSC cell line differentiation results. Top row shows FACS scatterplots showing a lower percentage of CD31/PDPN double positive cells when treated with no sodium acetate, and the bottom row shows the increase in the double positive population with the sodium acetate treatment.

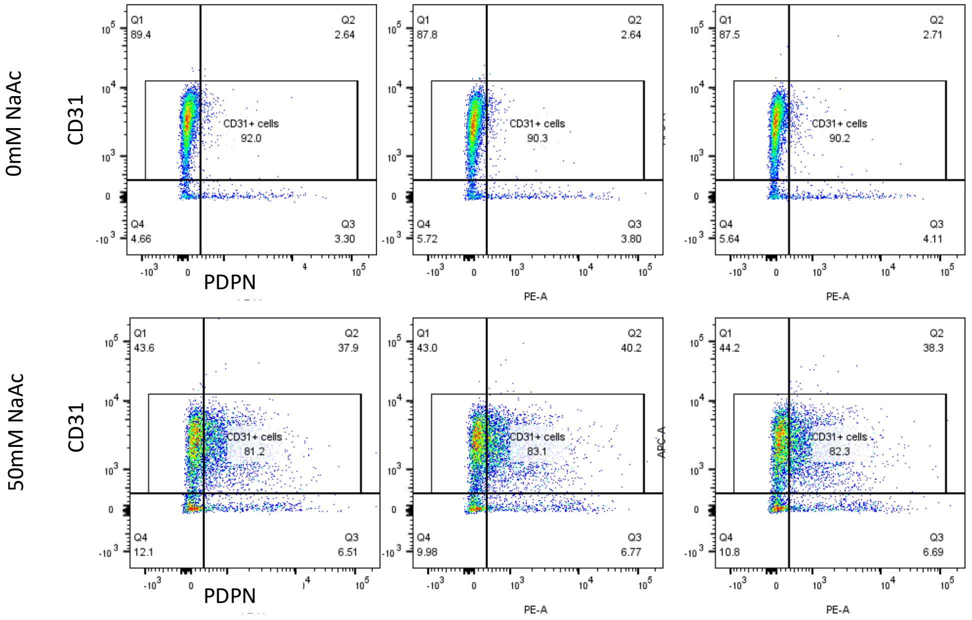


**Supplementary Figure 4**. Human 1019 iPSC cell line differentiation results. Top row shows FACS scatterplots showing a lower percentage of CD31/PDPN double positive cells when treated with no sodium acetate, and the bottom row shows the increase in the double positive population with the sodium acetate treatment.

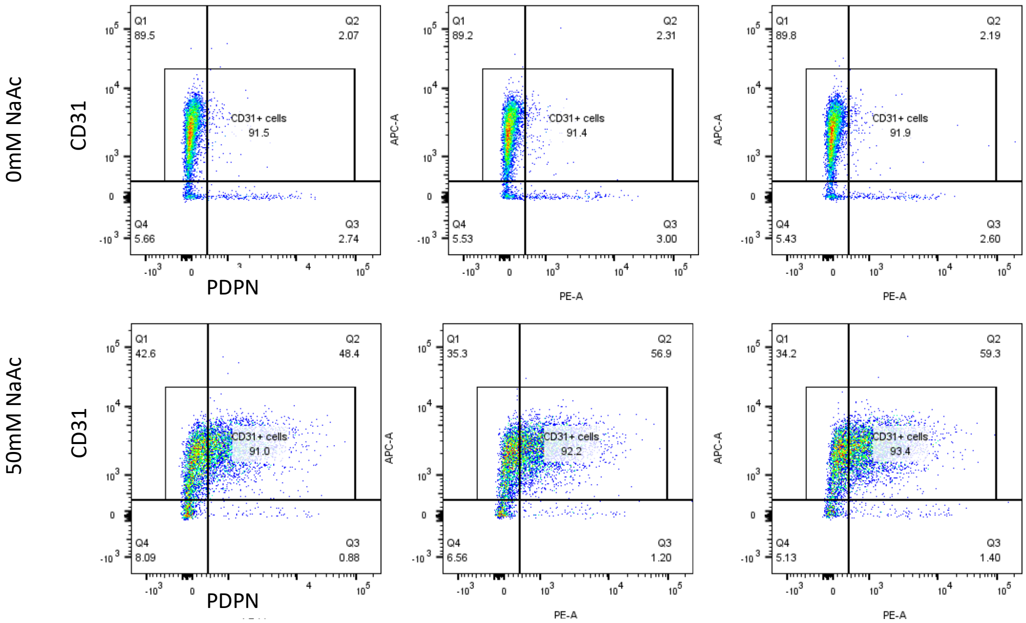


**Supplementary Figure 5.** Human 5807 iPSC cell line differentiation results. Top row shows FACS scatterplots showing a lower percentage of CD31/PDPN double positive cells when treated with no sodium acetate, and the bottom row shows the increase in the double positive population with the sodium acetate treatment.

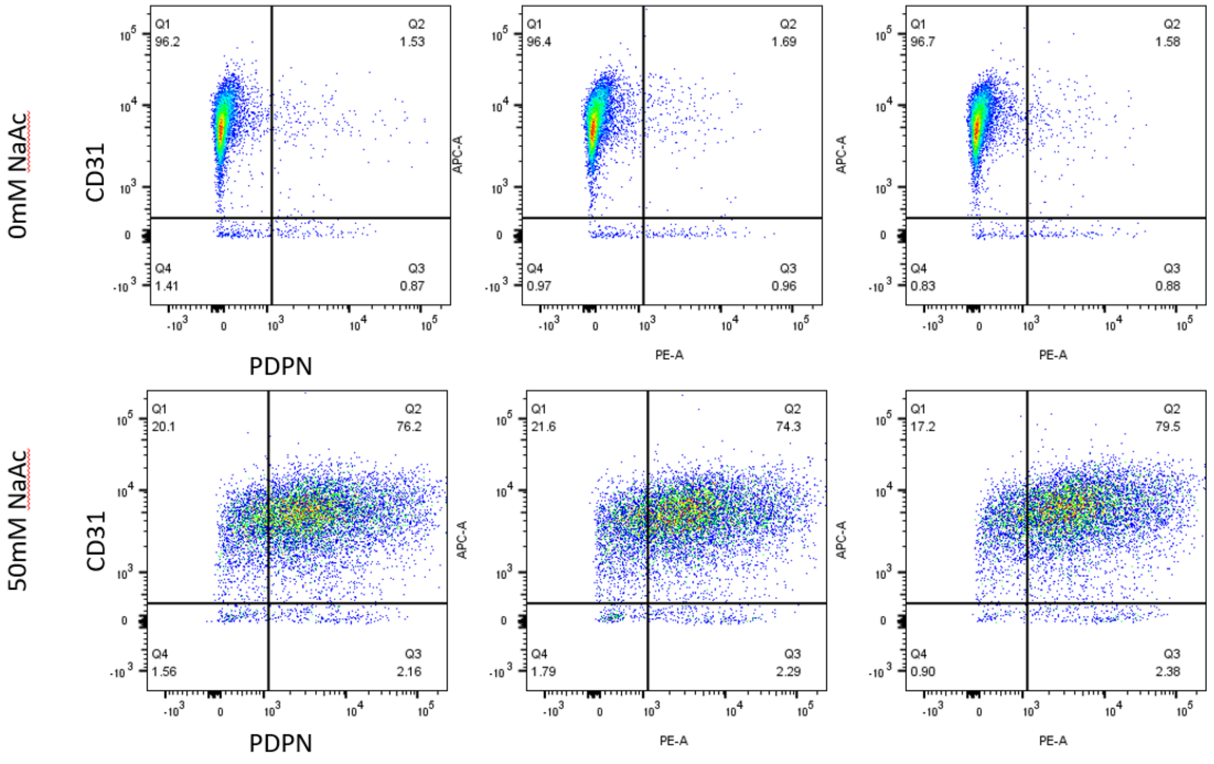


**Supplementary Figure 6.** Human IMR90 iPSC cell line differentiation results. Top row shows FACS scatterplots showing a lower percentage of CD31/PDPN double positive cells when treated with no sodium acetate, and the bottom row shows the increase in the double positive population with the sodium acetate treatment.

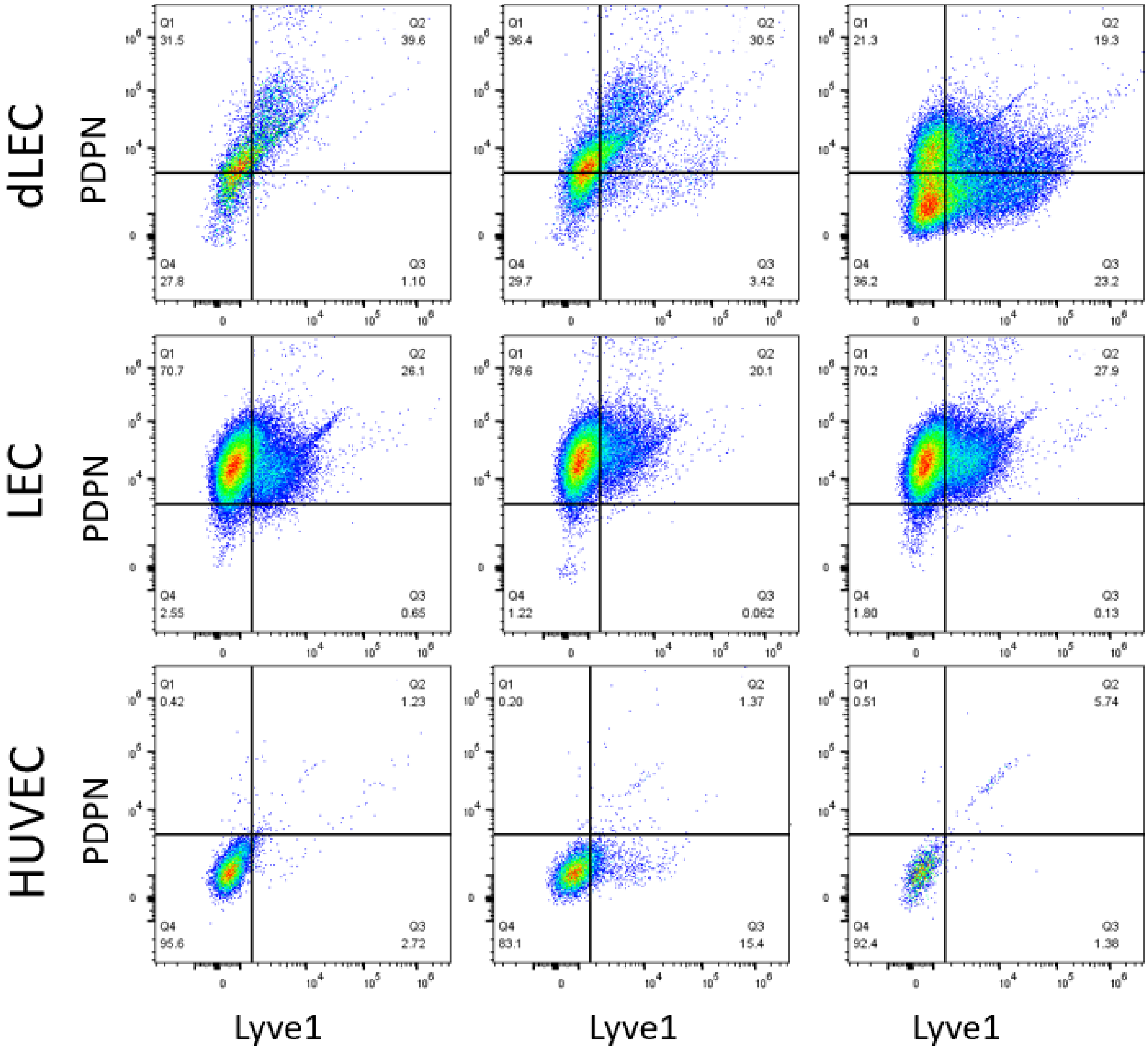


**Supplementary Figure 7.** Lyve1 and PDPN-positive populations for human IMR90-differentiated LEC compared to native human LEC (second row) and native human blood endothelial cell (third row).

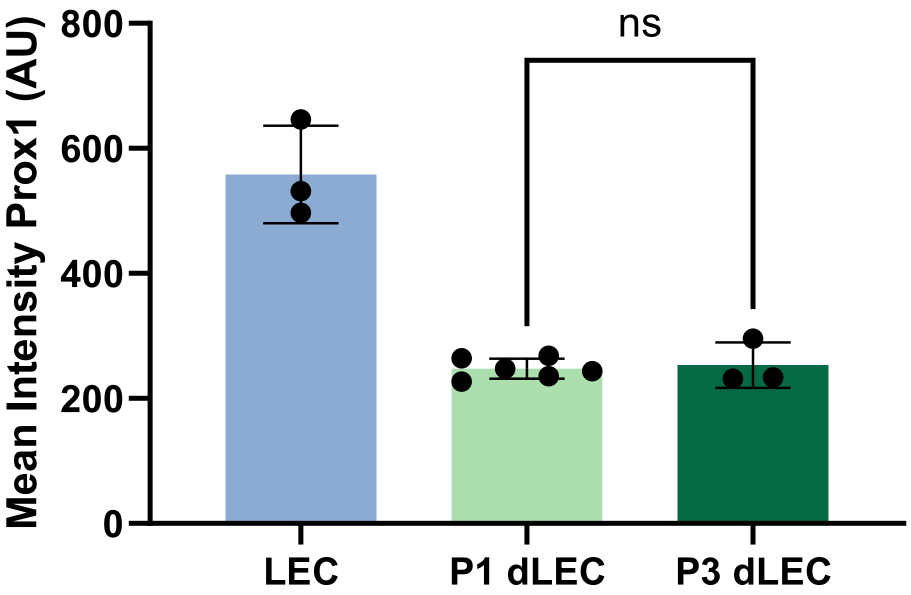


**Supplementary Figure 8.** Average Prox1 expression levels quantified from immunostaining data of IMR90-dLEC showing a non-significant change from P1 to P3.

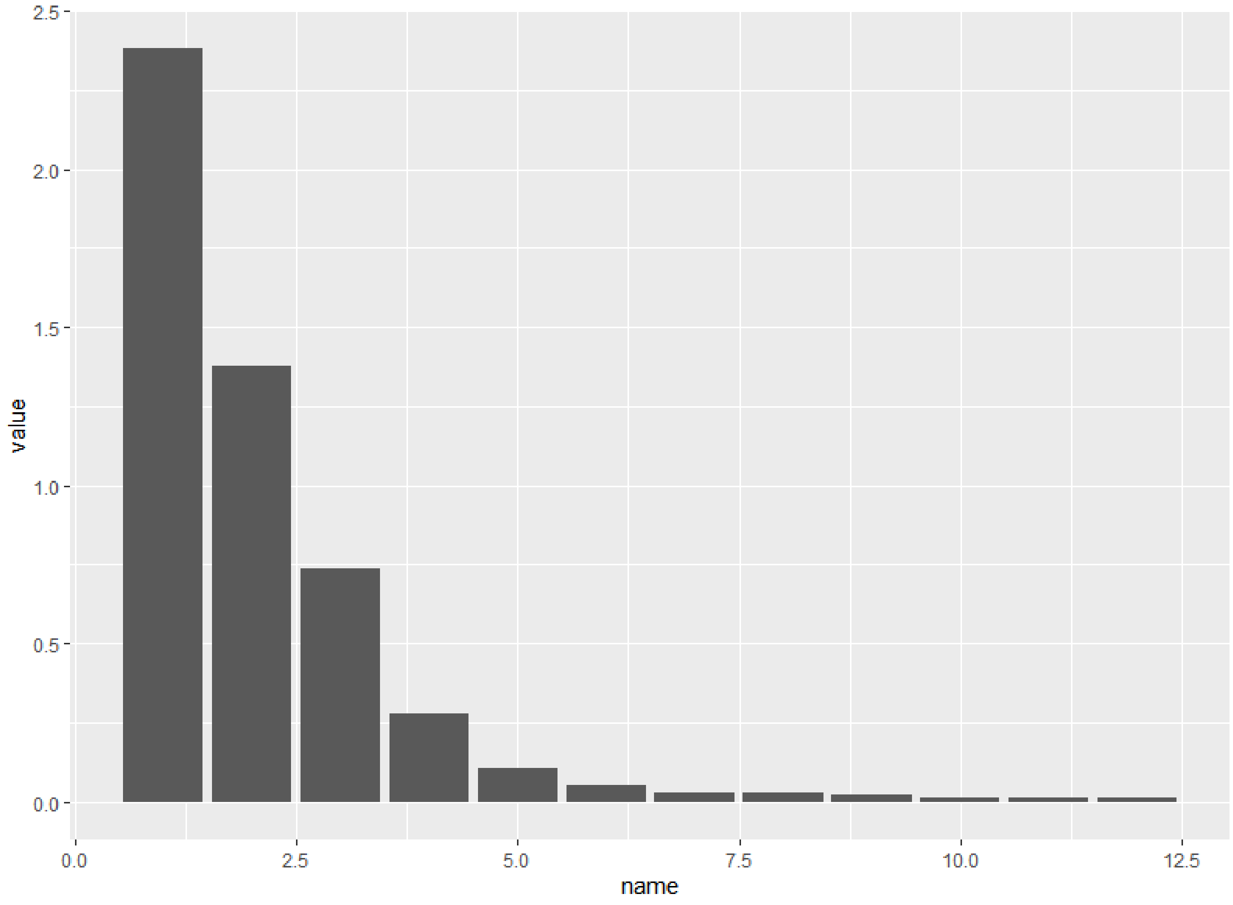


**Supplementary Figure 9**. Covariances for the principal component analysis. 74% of the variance is explained by the first two principal components.

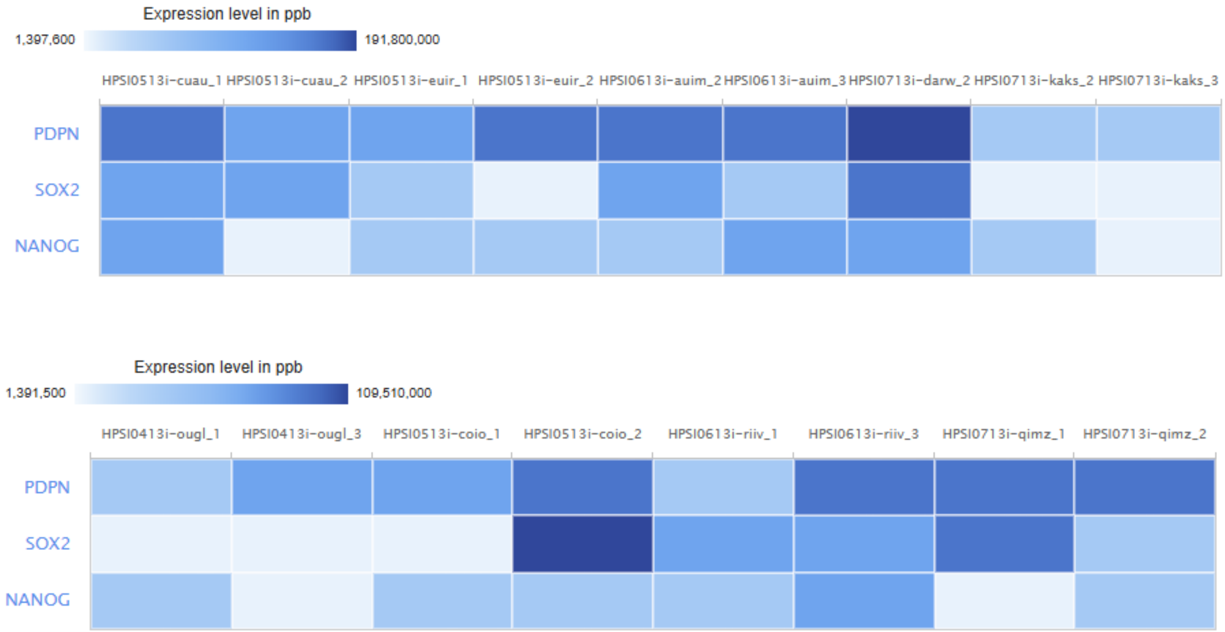


**Supplementary Figure 10.** Expression levels of PDPN from the proteomics data provided by the Human Induced Pluripotent Stem Cells Initiative. The PDPN expression is compared with Sox2 and NANOG expressions, which are well-known pluripotency markers. The plots were generated using the user interface of the Expression Atlas through the European Bioinformatics Initiative website.

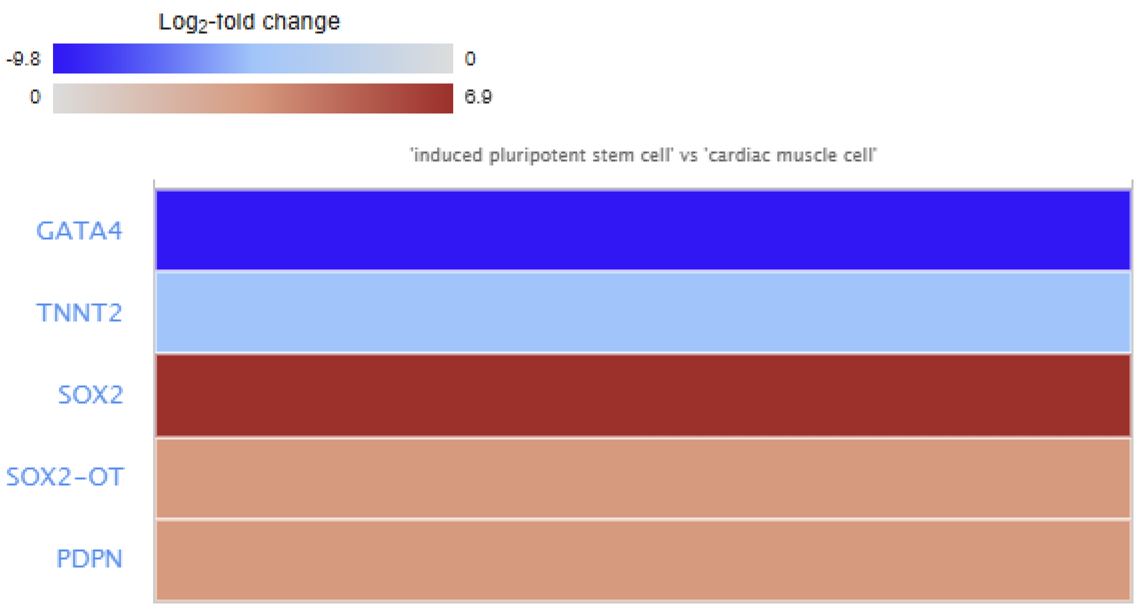


**Supplementary Figure 11.** Relative gene expression levels of iPSCs compared to iPSC-derived cardiomyocytes. TNNT2 and GATA4, well-known cardiomyocyte markers, are highly upregulated in the cardiomyocytes, whereas Sox2 is downregulated. PDPN is also similarly downregulated in differentiated cardiomyocytes, indicating that iPSCs initially express PDPN and subsequently lose the PDPN expression during the differentiation process. The figures were generated using the web interface of the European Bioinformatics Initiative. The raw data was provided by Banovich and colleagues in their 2018 publication.

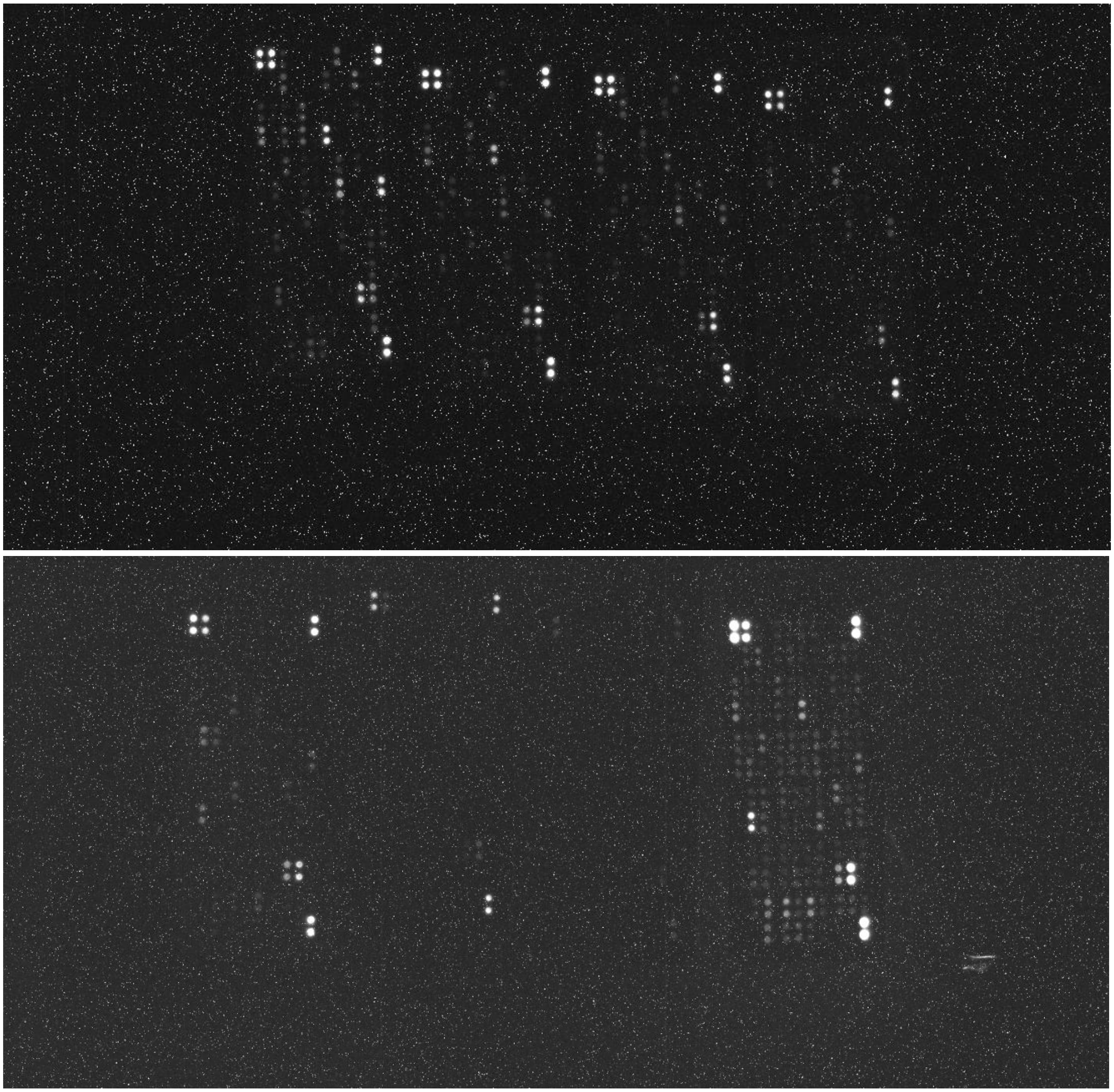

**Supplementary Figure 12**. Raw images of the cytokine assay kit. First image is of the angiogenesis assay kit and the second image is of the inflammatory cytokine assay kit.

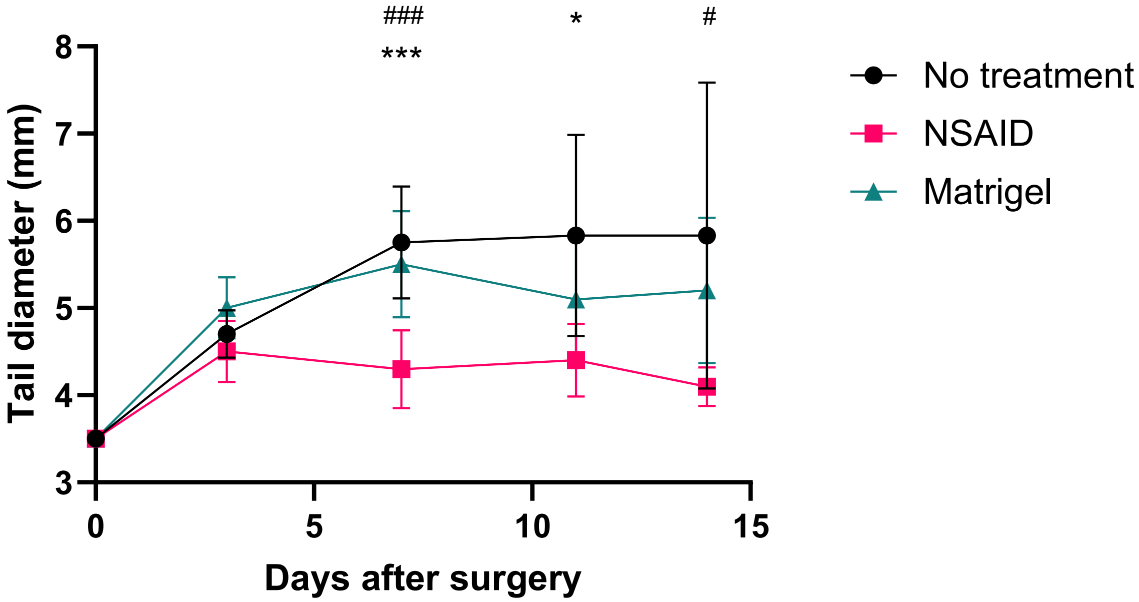


$$d = \frac{\mu_{NT} - \mu_{NSAID}}{stdev} \approx 2.25$$

For $\alpha = 0.05, \beta = 0.8, \; n = (Z_{1-\frac{\alpha}{2}} + Z_{1-\beta})^2 * \frac{2}{d^2} \approx 3.1$

**Supplementary Figure 13.** Pilot study results showing the difference in degree of swelling for NSAID and control groups. The results from day 7, which was decided as the endpoint for subsequent studies, were used to perform power analysis. Student t-test was performed between each sample type at each timepoint. * indicates significance between No Treatment and NSAID group, and # between No treatment and Matrigel group. * or # p<0.05, ***or ### p<0.001. Subsequent power analysis was performed between the negative control (No Treatment) and positive control (NSAID). Alpha value of 0.05 and beta value of 0.8 were selected.

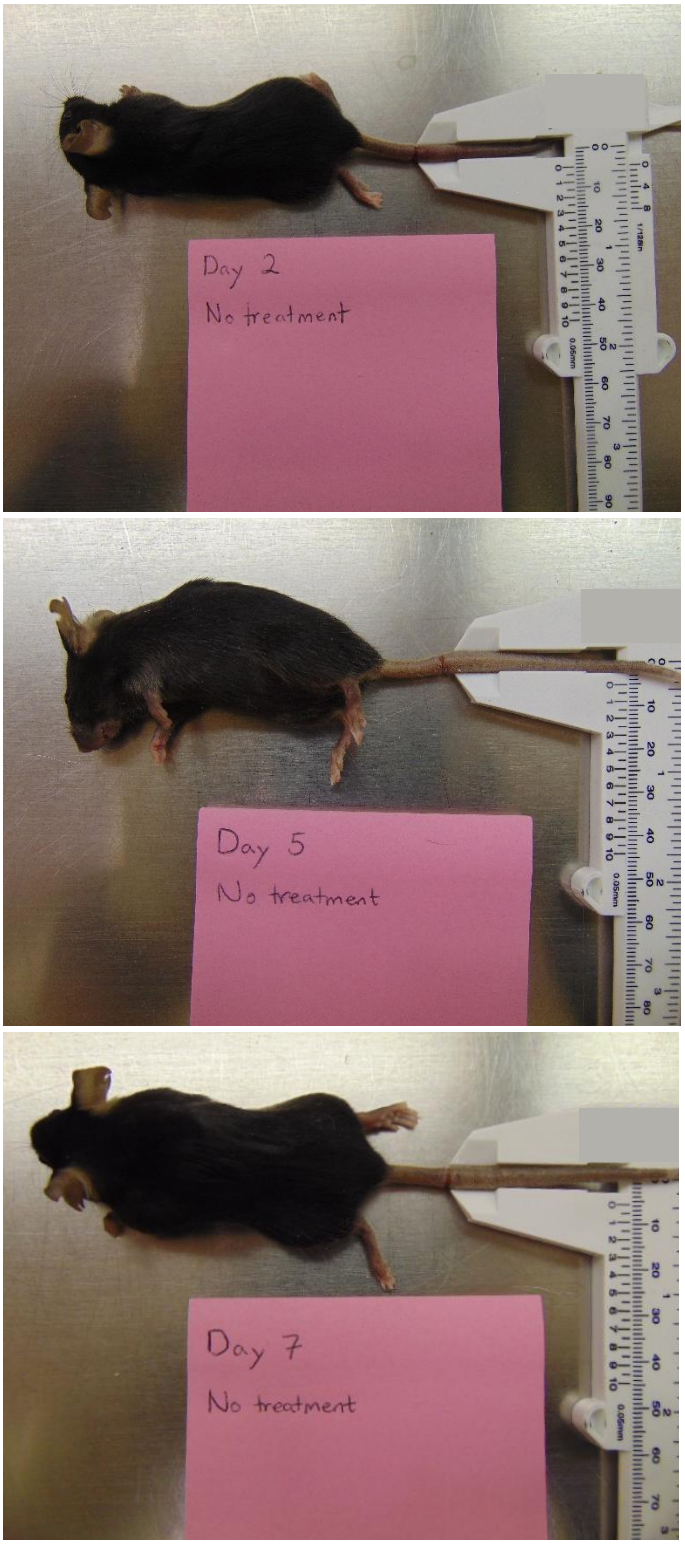


**Supplementary Figure 14.** Representative images of mice in the no treatment group at day 2, 5, and 7.

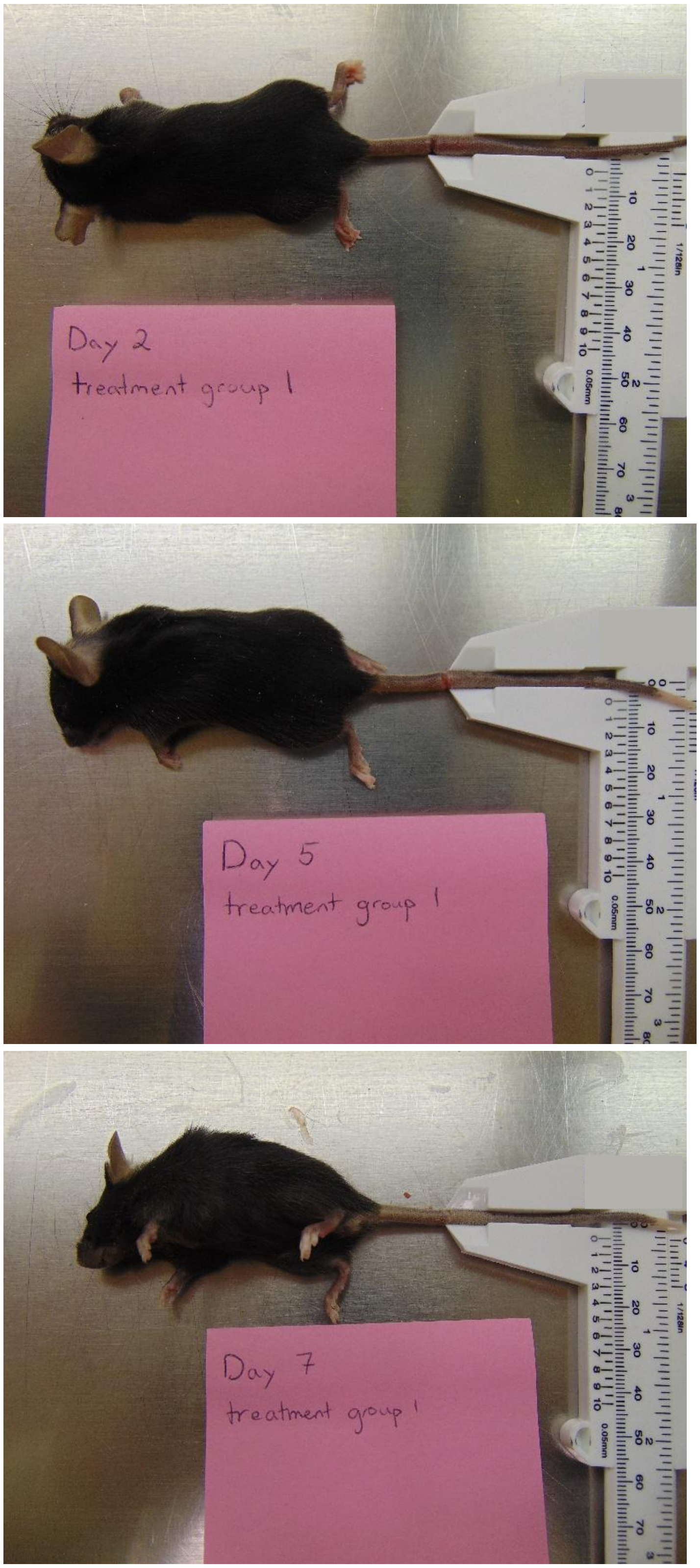


**Supplementary Figure 15.** Representative images of mice in the dLEC treatment group at day 2, 5, and 7.

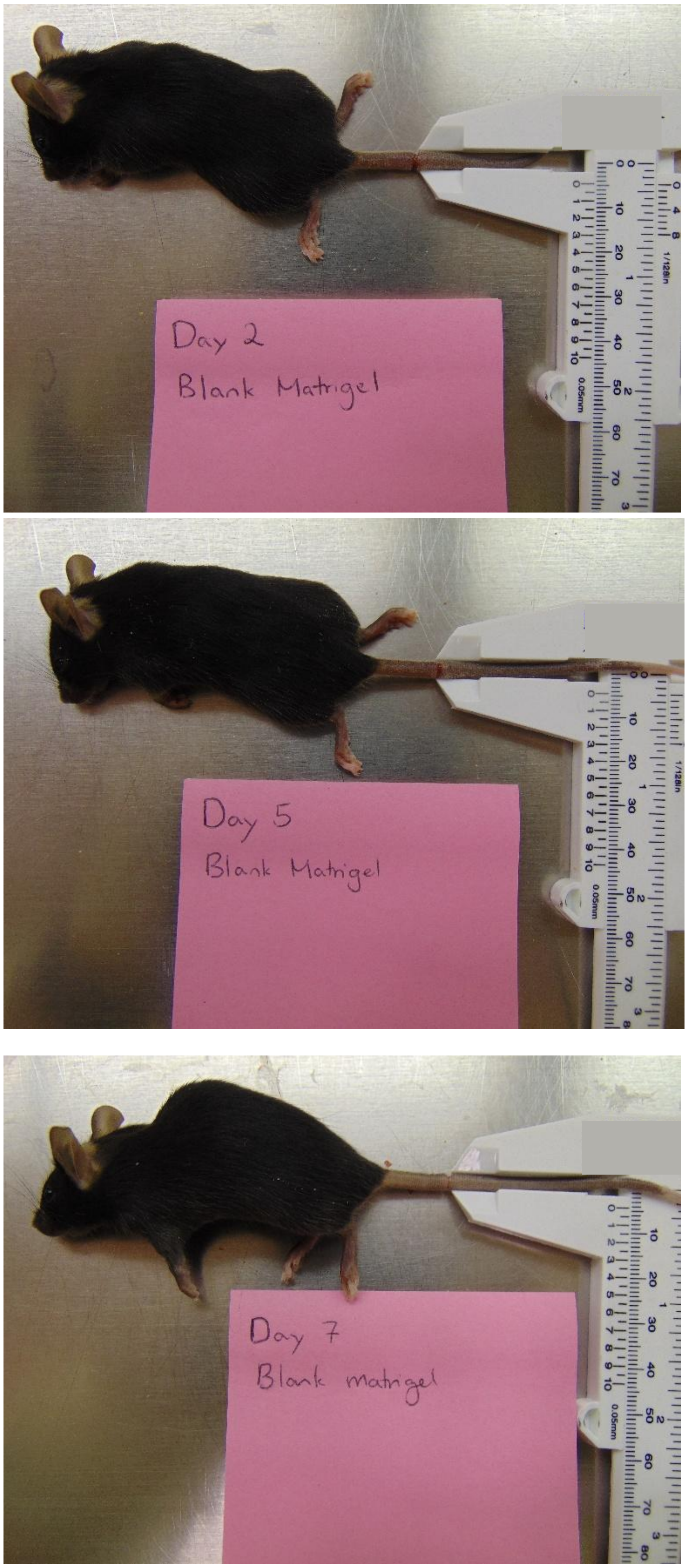


**Supplementary Figure 16**. Representative images of mice in the blank Matrigel treatment group at day 2, 5, and 7.

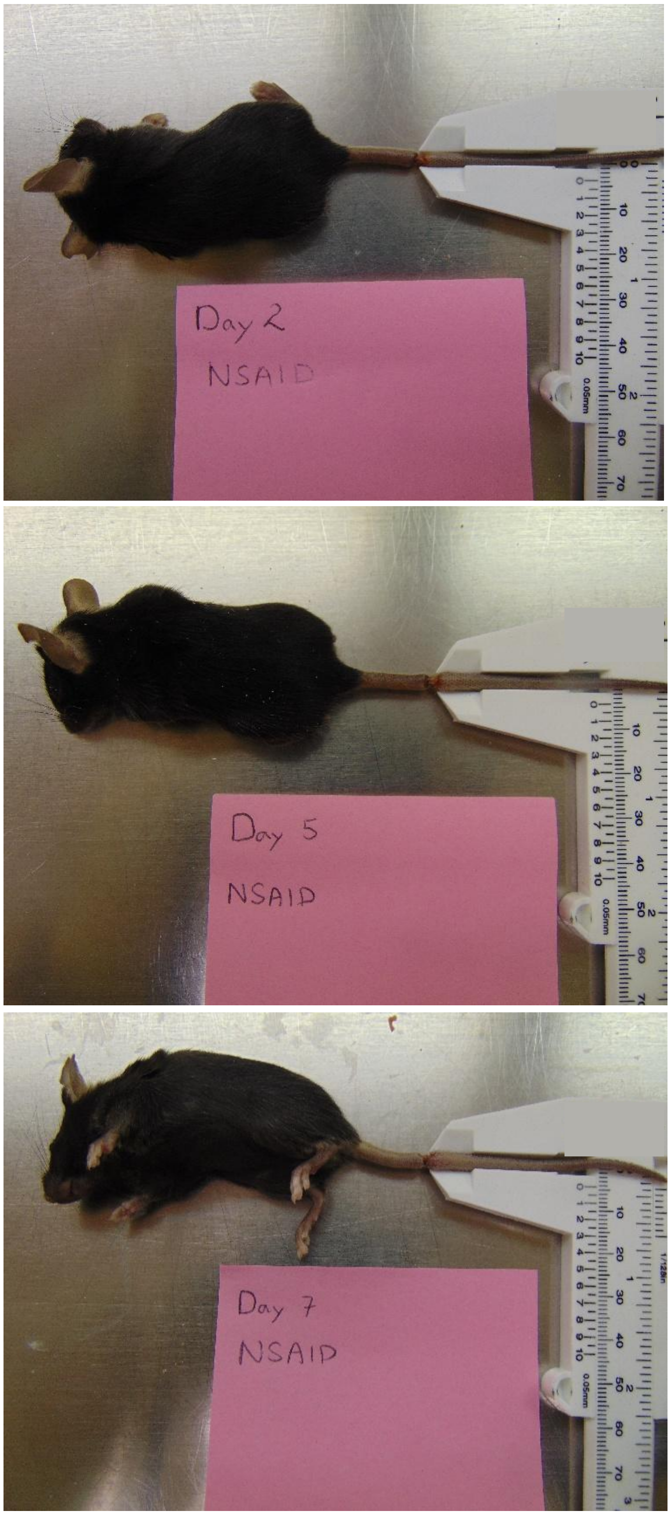


**Supplementary Figure 17.** Representative images of mice in the NSAID treatment group at day 2, 5, and 7.

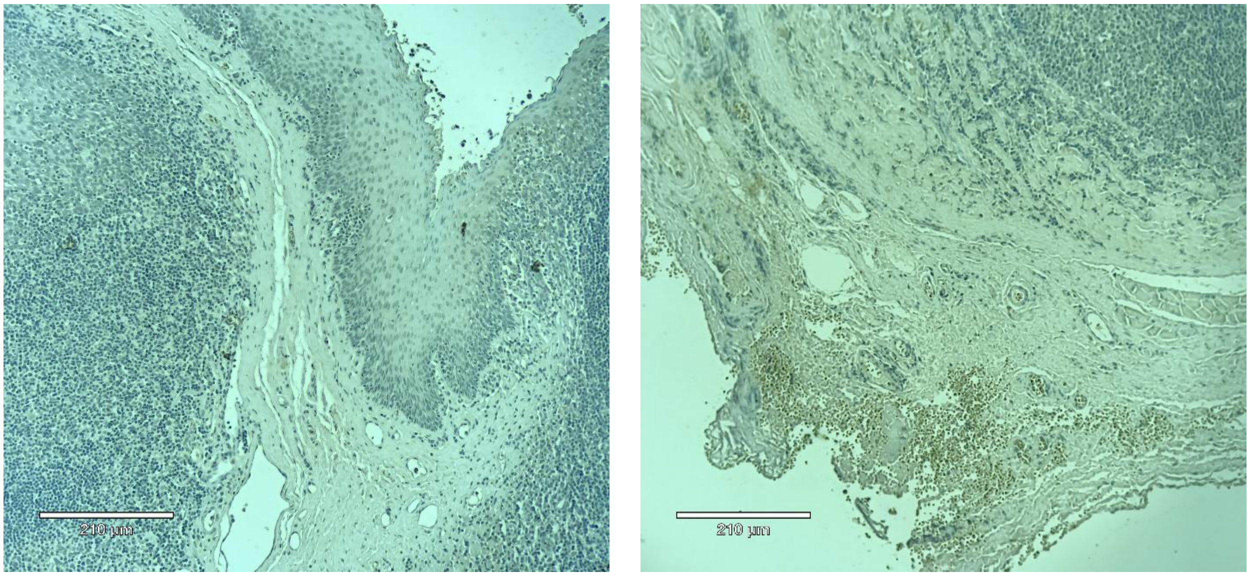


**Supplementary Figure 18**. IgG control staining for mouse tail lymphedema model samples showing a lack of positive brown stains. Left is anti-hamster IgG and right is anti-mouse IgG.

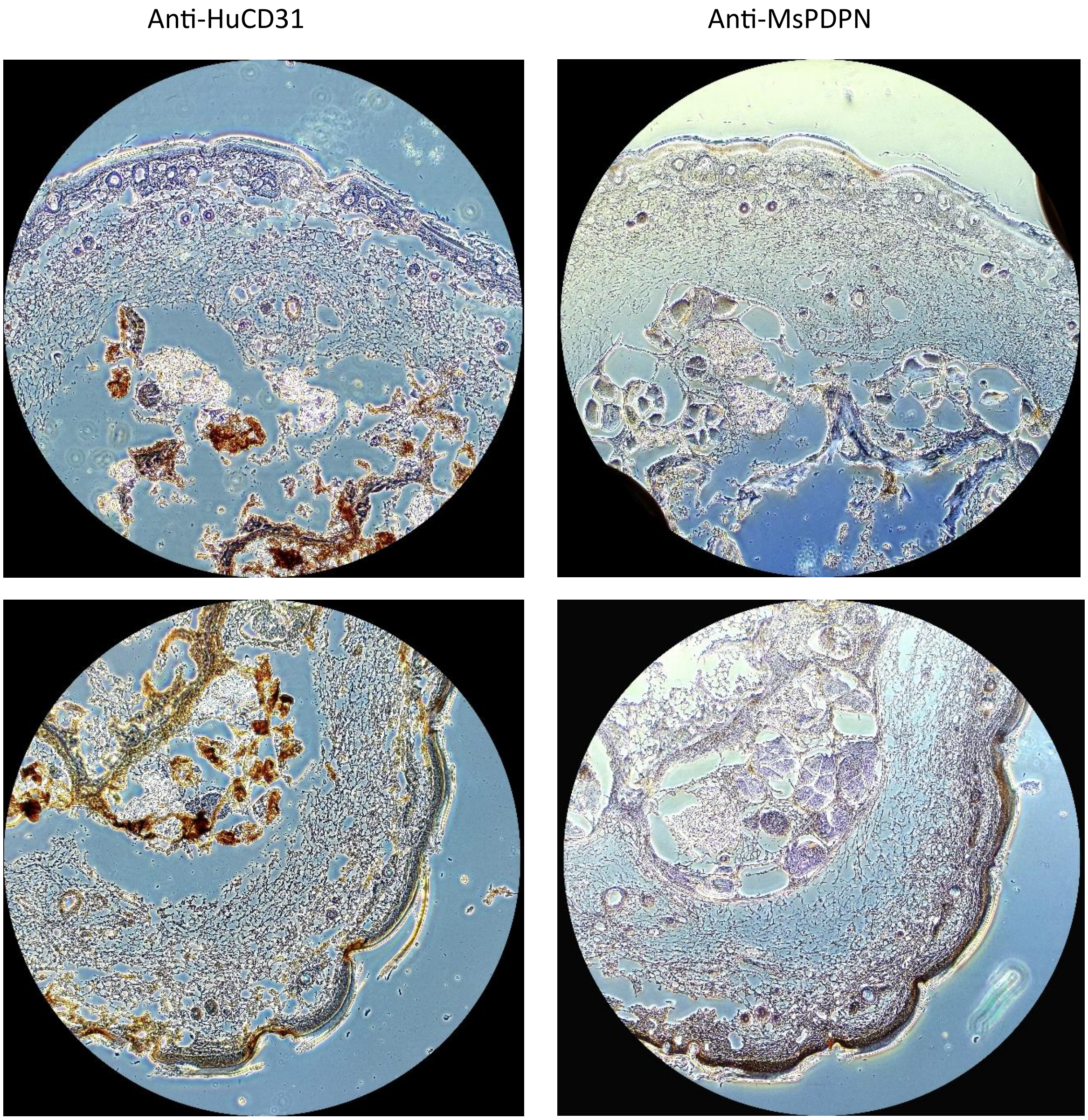


**Supplementary Figure 19**. Representative images of lymphatic vessels for blank Matrigel-treated mice.

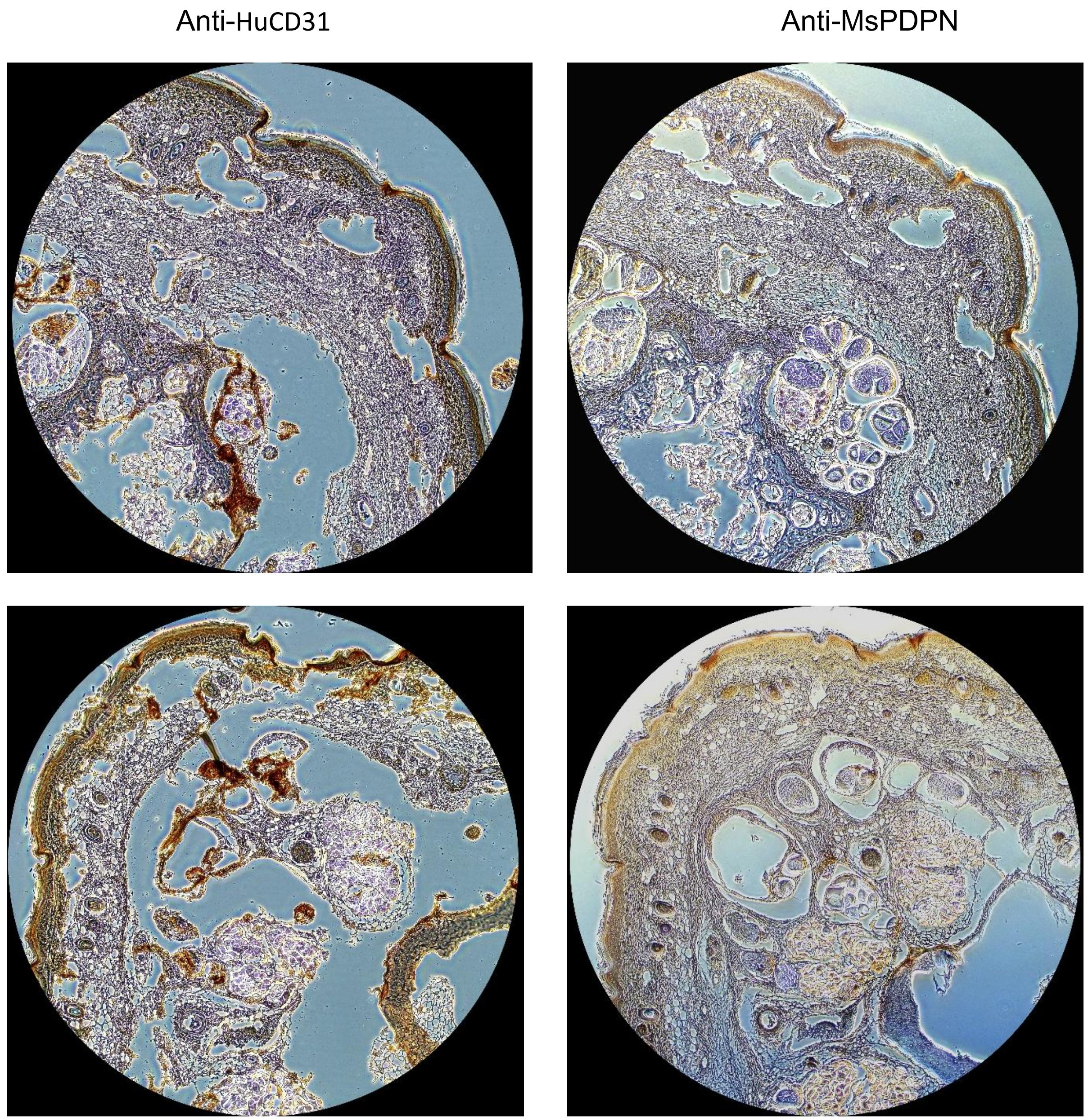


**Supplementary Figure 20**. Representative images of lymphatic vessels for Matrigel-dLEC treated mice.

| Cell line | Generator | Sex | Source | Reprogramming |
|---|---|---|---|---|
| i026 | Klinikum der Universität zu Köln | M | Peripheral blood mononuclear cell | Sendai virus |
| 5807 | WiCell | F | Skin fibroblast | Episomal plasmid |
| IMR90 | WiCell | F | Lung fibroblast | Lentiviral transfection |
| 1019 | ATCC | M | Foreskin fibroblast | Sendai virus |

**Supplementary Table 1.** List of iPSC cell lines used in differentiation.

| Gene (Entrez Gene ID) | Reporter type | Taqman Probe ID (Thermo Fisher) |
|---|---|---|
| **Human PDPN (10630)** | FAM-MGB | Hs00366766_m1 |
| **Human Flt4 (2324)** | FAM-MGB | Hs01047677_m1 |
| **Human Prox1 (5629)** | FAM-MGB | Hs00896293_m1 |
| **Human Cdh4 (1002)** | FAM-MGB | Hs00899698_m1 |
| **Human GAPDH (2597)** | FAM-MGB | Hs99999905_m1 |

**Supplementary Table 2.** List of RT-qPCR Taqman primers.

| Target/Conjugate | Host / Isotype | Application | Dilution | Vendor |
|---|---|---|---|---|
| **Human Prox1 - Unconjugated** | Goat IgG | Immunostaining | 1:200 | Abcam |
| **Human Prox1 - FITC** | Mouse IgG1 | Flow Cytometry | 1:200 | Novus Biologicals |
| **Human PDPN - Unconjugated** | Mouse IgG1 | Immunostaining | 2μg/mL | Abcam |
| **Human PDPN - PE** | Hamster IgG | Flow Cytometry | 1:400 | Novus Biologicals |
| **Human ERG - Unconjugated** | Rabbit IgG | Immunostaining | 1:200 | Abcam |
| **Human ERG - PE** | Mouse IgG1 | Flow Cytometry | 1:200 | Novus Biologicals |
| **Human CD31 - APC** | Mouse IgG1 | Flow Cytometry | 1:100 | Bio-Techne |
| **Human Lyve1 - APC** | Rabbit IgG | Flow Cytometry | 1:200 | Novus Biologicals |
| **Goat IgG - AF488** | Donkey IgG | Immunostaining | 1:500 | Abcam |
| **Rabbit IgG - AF594** | Goat IgG | Immunostaining | 1:500 | Abcam |
| **Mouse IgG - AF647** | Donkey IgG | Immunostaining | 1:500 | Abcam |
| **Human VECad - AF647** | Mouse IgG | Hydrogel staining | 1:50 | Santa Cruz Biotechnologies |
| **Human PDPN - AF488** | Mouse IgG2 | Hydrogel Staining | 1:50 | Santa Cruz Biotechnologies |
| **Human CD31 - Unconjugated** | Mouse IgG | IHC | 1:50 | Dako |
| **Mouse PDPN - Unconjugated** | Hamster IgG | IHC | 1:50 | Thermo Fisher |
| **Mouse IgG - HRP** | Goat IgG | IHC | 1:200 | Santa Cruz Biotechnologies |
| **Hamster IgG - HRP** | Goat IgG | IHC | 1:200 | Santa Cruz Biotechnologies |

**Supplementary Table 3.** List of antibodies used for immunostaining, IHC, and FACS analysis.

| Cell line | Vendor | Sex | Source | Catalog |
|---|---|---|---|---|
| BEC | Promocell | M | Human dermal blood endothelial cells, juvenile foreskin | C-12225 |
| LEC | Promocell | M | Human dermal lymphatic endothelial cells, juvenile foreskin | C-12217 |
| HUVEC | Promocell | F | Human umbilical vein endothelial cells | C-12200 |

**Supplementary Table 4.** Primary human cell lines used in this study.
